# Genome-Wide Scan for Adaptive Divergence and Association with Population-Specific Covariates

**DOI:** 10.1101/023721

**Authors:** Mathieu Gautier

## Abstract

In population genomics studies, accounting for the neutral covariance structure across population allele frequencies is critical to improve the robustness of genome-wide scan approaches. Elaborating on the BayEnv model, this study investigates several modeling extensions i) to improve the estimation accuracy of the population covariance matrix and all the related measures; ii) to identify significantly overly differentiated SNPs based on a calibration procedure of the XtX statistics; and iii) to consider alternative covariate models for analyses of association with population-specific covariables. In particular, the auxiliary variable model allows to deal with multiple testing issues and, providing the relative marker positions are available, to capture some Linkage Disequilibrium information. A comprehensive simulation study was carried out to evaluate the performances of these different models. Also, when compared in terms of power, robustness and computational efficiency, to five other state-of-the-art genome scan methods (BayEnv2, BayScEnv, BayScan, FLK and LFMM) the proposed approaches proved highly effective. For illustration purpose, genotyping data on 18 French cattle breeds were analyzed leading to the identification of thirteen strong signatures of selection. Among these, four (surrounding the KITLG, KIT, EDN3 and ALB genes) contained SNPs strongly associated with the piebald coloration pattern while a fifth (surrounding PLAG1) could be associated to morphological differences across the populations. Finally, analysis of Pool–Seq data from 12 populations of *Littorina saxatilis* living in two different ecotypes illustrates how the proposed framework might help addressing relevant ecological issue in non–model species. Overall, the proposed methods define a robust Bayesian framework to characterize adaptive genetic differentiation across populations. The BayPass program implementing the different models is available at http://www1.montpellier.inra.fr/CBGP/software/baypass/.

---

Contrasting patterns of local genetic variation over the whole genome represents a valuable strategy to identify loci underlying the response to adaptive constraints (Cavalli-Sforza 1966). As further noted by Lewontin and Krakauer (1973): “*while natural selection will operate differently for each locus and each allele at a locus, the effect of breeding structure is uniform over all loci and all alleles”.* Hence, genome scan approaches to detect footprints of selection aim at discriminating among the global effect of the demographic evolutionary forces (e.g., gene flow, inbreeding and genetic drift) from the local effect of selection (Vitalis *et al.* 2001; Balding and Nichols 1995). In practice, applications of these methods have long been hindered by technical difficulties in assessing patterns of genetic variation on a whole genome scale. However, the advent of next-generation sequencing and genotyping molecular technologies now allows to provide a detailed picture of the structuring of genetic variation across populations in both model and non–model species (Davey *et al.* 2011). As a result, in the population genomics era, a wide range of approaches have been developed and applied to detect selective sweeps using population data (see Vitti *et al.* 2013; Oleksyk *et al.* 2010, for reviews). Among these, population differentiation (*F*_ST_) based methods still remain among the most popular, particularly in non–model species since they do not require accurate genomic resources (e.g., physical or linkage maps) and experimental designs with only a few tens of genotyped individuals per population are generally informative enough. Also, *F*_ST_–based methods are well suited to the analysis of data from Pool–Seq experiments that consist in sequencing pools of individual DNAs (Schlötterer *et al.* 2014) and provide cost-effective alternatives to facilitate and even improve allele frequency estimation at genome-wide markers (Gautier *et al.* 2013).

In practice, assuming the vast majority of the genotyped markers behave neutrally, overly differentiated loci that are presumably subjected to selection might simply be identified from the extreme tail of the empirical distribution of the locus–specific *F*_ST_ (Akey *et al.* 2002; Weir *et al.* 2005; Flori *et al.* 2009). Even if such a model–free strategy does not rely on any arbitrary assumptions about the (unknown) demographic history of the sampled populations, it prevents from controlling for false positive (and negative) signals. Conversely, model–based approaches have also been developed and are basically conceived as locus–specific tests of departure from expectation under neutral demographic models (e.g, Gautier *et al.* 2010a). These include, for instance, demographic models under pure-drift (Gautier *et al.* 2010a; Nicholson *et al.* 2002) and at migration-drift equilibrium without (Beaumont and Balding 2004; Foll and Gaggiotti 2008; Riebler *et al.* 2008; Guo *et al.* 2009) or with selection (Vitalis *et al.* 2014). Although robust, to some extent, to more complex history (Beaumont and Nichols 1996; Beaumont 2005), these methods remain limited by the oversimplification of the underlying demographic models. In particular, hierarchically structured population history, as produced under tree–shaped phylogenies, have been shown to increase false positive rates (Excoffier *et al.* 2009). To cope with these issues, two kinds of modeling extensions have recently been explored. They either rely on hierarchical island models, thus requiring a prior definition of the sampled population relationships (Foll *et al.* 2014; Gompert *et al.* 2010), or consist in estimating the correlation structure of allele frequencies across the populations that originates from their shared history (Bonhomme *et al.* 2010; Coop *et al.* 2010; Günther and Coop 2013).

Whatever the method used, the main limitation of the indirect genome scan approaches ultimately resides in the biological interpretation of the footprints of selection identified, i.e., to which adaptive constraints the outlier loci are responding. In species with functionally annotated reference genomes, the characterization of co-functional relationships among the genes localized within regions under selection might help gaining insights into the underlying driving physiological pathways (e.g., Flori *et al.* 2009). Although, following a “reverse ecology” approach (Li *et al.* 2008), they may further lead to the definition of candidate adaptive traits for validation studies, such interpretations remain prone to misleading storytelling issues (Pavlidis *et al.* 2012). Alternatively, prior knowledge about some characteristics discriminating the populations under study could provide valuable insights. Focusing on environmental gradients, several approaches have recently been proposed to evaluate association of ecological variables with marker genetic differentiation by extending *F*_ST_–based models (Coop *et al.* 2010; de Villemereuil and Gaggiotti 2015; Frichot *et al.* 2013; Günther and Coop 2013; Guillot *et al.* 2014; Hancock *et al.* 2011, 2008; Joost *et al.* 2007; Poncet *et al.* 2010). The rationale is that environmental variables distinguishing the differentiated populations should be associated with allele frequencies differences at loci subjected to the selective constraints they impose (Coop *et al.* 2010). In principle, such population-based association studies may also be more broadly relevant to any quantitative or categorical population–specific covariable. More generally, as for the covariable-free genome–scan approaches, accounting for the neutral correlation of allele frequencies across populations is critical for these methods (de Villemereuil *et al.* 2014; De Mita *et al.* 2013).

Overall, the Bayesian hierarchical model proposed by Coop *et al.* (2010) and implemented in the BayEnv2 software represents a flexible framework to address these issues. It indeed allows to both identify outlier loci (Günther and Coop 2013) and to further annotate the resulting footprints of selection by quantifying their association with population-specific covariables (if available). A key parameter of the model is the (scaled) population covariance matrix across population allele frequencies. Although this matrix might be viewed as purely instrumental, it explicitly incorporates their neutral correlation structure and is in turn highly informative for demographic inference purposes (Lipson *et al.* 2013; Pickrell and Pritchard 2012). Elaborating on the BayEnv model (Coop *et al.* 2010; Günther and Coop 2013), the purpose of this paper is threefold. First, we introduce modeling modifications and extensions to improve the estimation accuracy of the population covariance matrix and the different related measures. Second, we propose a posterior checking procedure to identify markers subjected to adaptive differentiation based on a calibration of the XtX statistics (Günther and Coop 2013). Third, we investigate alternative modeling strategies and decision criteria to perform association studies with population–specific covariables. In particular, we introduce a model with a binary auxiliary variable to classify each locus as associated or not. Through the prior distribution on this latter variable, the approach deals with the problem of multiple testing (e.g. Riebler *et al.* 2008). In addition, providing information about marker positions is available, this modeling also allows to account for Linkage Disequilibrium (LD) between markers via an Ising prior. As a by–product of this study, a user–friendly and freely available program, named BayPass, was developed to implement inferences under the different models. To evaluate the accuracy of the methods, we further carry out comprehensive simulation studies. In addition, two real data sets were analyzed in more detail to illustrate the range of application of the methods. The first consists of 453 individuals from 18 French cattle breeds genotyped at 42,056 SNPs (Gautier *et al.* 2010b) and the second of a Pool–Seq data on 12 *Littorina saxatilis* populations from three distinct geographical regions and living in two different ecotypes (Westram *et al.* 2014).

## Models

In the following we describe the different Bayesian hierarchical models considered in this study and implemented in the BayPass program. Consider a sample made of *J* populations (sharing a common history) with a label, *j*, which varies from 1 to *J*. The data consist of *I* SNP loci, which are biallelic markers with reference allele arbitrarily defined (e.g., by randomly drawing the ancestral or the derived state). Let *n_ij_* be the total number of genes sampled at the *j*^th^ locus (1 ≤ *i* ≤ *I*) in the *j*^th^ population (1 *≤ j ≤ I*), that is, twice the number of genotyped individuals in a diploid population. Let *y_ij_* be the count of the reference allele at the *i*^th^ locus in the *j*^th^ sampled population. When considering allele count data, the *y_ij_*’s (and the *n_ij_*’s) are the observations while for Pool–Seq data, read count are observed instead. In this case, the *n_ij_*’s correspond for all the markers within a given pool to its haploid sample size *n_j_* (i.e., twice the number of pooled individuals for diploid species). Let further *c_ij_* be the (observed) total number of reads and *r_ij_* the (observed) number of reads with the reference allele. For Pool–seq data, to integrate over the unobserved allele count, the conditional distribution of the *r_ij_* given *c_ij_,n_j_* and the (unknown) *y_ij_* is assumed binomial (Gautier *et al.* 2013; Günther and Coop 2013): 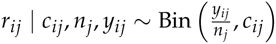.

Assuming Hardy–Weinberg Equilibrium, the conditional distribution of *y_ij_* given *n_ij_* and the (unknown) allele frequency *α_ij_* is also assumed binomial:

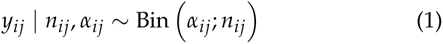

Note that this corresponds to the first level (likelihood) of the hierarchical model when dealing with allele count data and to the second level (prior) for Pool–Seq data. As previously proposed and discussed (Coop *et al.* 2010; Gautier *et al.* 2010a; Nicholson *et al.* 2002), for each SNP *i* and population *j* an instrumental variable 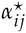 taking value on the real line is further introduced such that: 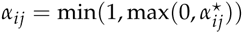. As represented in Figure 1, three different sub–classes of models are considered (each with their allele and read counts version). They are hereafter referred to as i) the core model (Figure 1A); ii) the standard covariate (STD) model (Figure 1B) and; iii) the auxiliary variable covariate (AUX) model (Figure 1C). Note that the core model is nested within the STD model which is itself nested within the AUX model.

**Figure 1.**
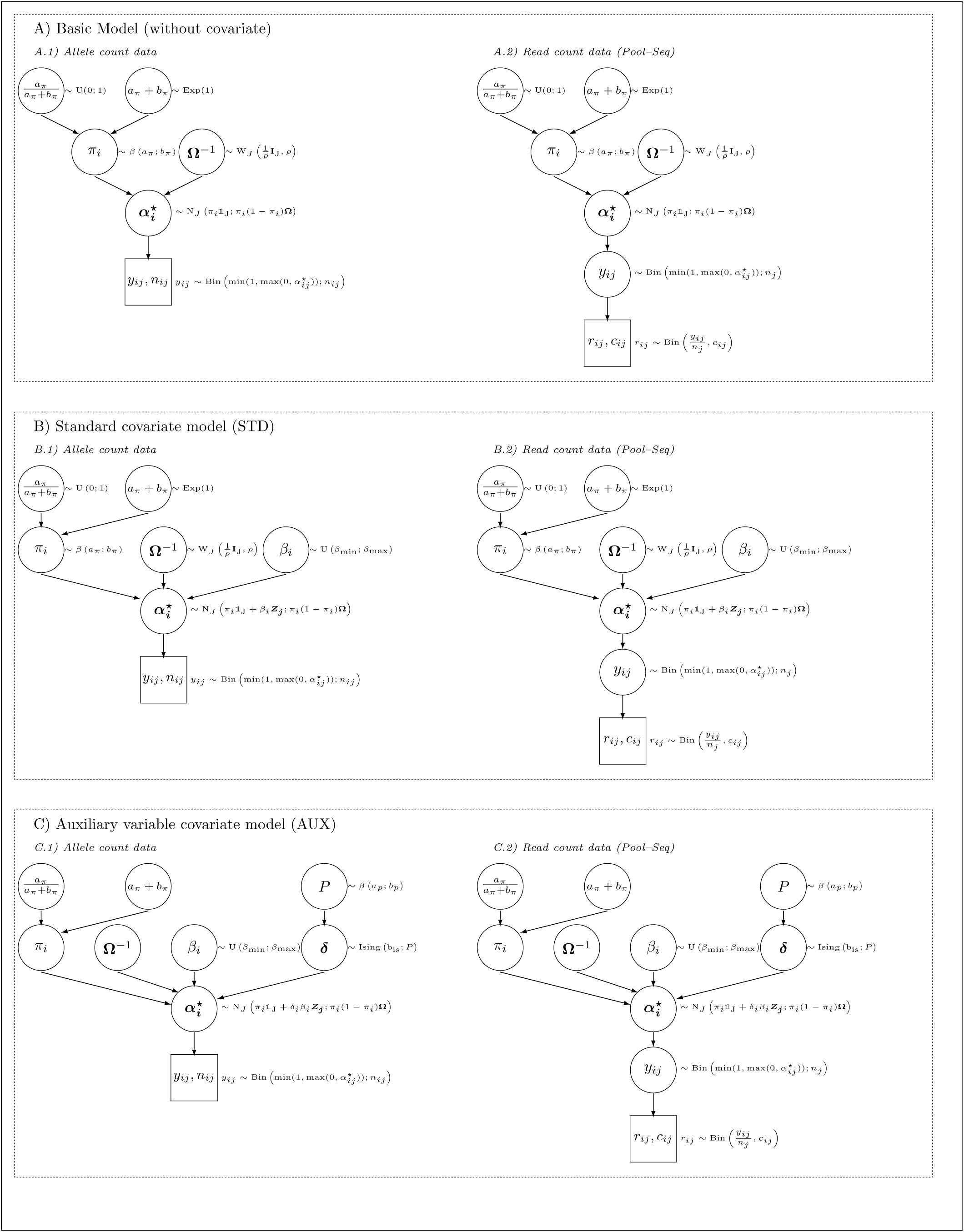
Directed Acyclic Graphs of the different hierarchical Bayesian models considered in the study and implemented in the BayPass software. See the main text for details about the underlying parameters and modeling assumptions.

### The core model

The core model (Figure 1A) is a multivariate generalization of the model by Nicholson *et al.* (2002) that was first proposed by Coop *et al.* (2010). For each SNP *i*, the prior distribution of the vector 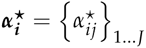 is multivariate Gaussian:

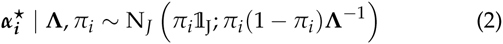
where 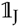 is a all-one vector of length *J*; the precision matrix Λ is the inverse of the (scaled) covariance matrix Ω (Λ = Ω^−1^) of the population allele frequencies; and *π_i_* is the weighted mean reference allele frequency that might be interpreted as the ancestral population allele frequency (Coop *et al.* 2010; Pickrell and Pritchard 2012). The *π_i_* are assumed Beta distributed:

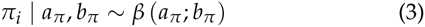

In such models, the parameters *a_π_* and *b_π_* are frequently fixed. For instance in BayEnv2 (Coop *et al.* 2010), *a_π_ = b_π_* = 1 leading to a uniform prior on *π_i_* over the (0,1) support. However, these parameters may be easily estimated from the model by specifying a prior distribution on the mean 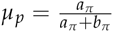 and the so-called “sample size” *v_p_ = a_π_* + *b_π_* (Kruschke 2014). Hence, a uniform and an exponential prior distribution are respectively considered for these two parameters:

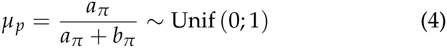
and

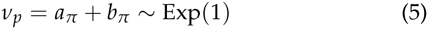

Finally, a Wishart prior distribution is assumed for the the precision matrix **Λ**:

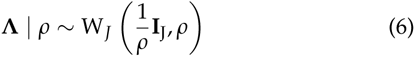
i.e., 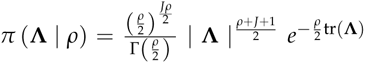 (I_J_ being the identity matrix of size *J*). For *ρ* ≥ *J* this is strictly equivalent to the parametrization introduced in Coop *et al.* (2010) who eventually came to fix *ρ* = *J.* Here, weaker informative priors are also explored with 0 < *ρ < J* (e.g., Gelman *et al.* 2003, p. 581) leading to so-called singular Wishart distributions. As will become apparent, *ρ* = 1 appears as the best default choice. Note however that inspection of the full conditional distribution of Λ (see File S1) suggests the influence of the prior might become negligible with increasing number of SNPs *I* and populations *J*.

### The standard covariate model (STD model)

The STD model represented in Figure 1B extends the core model as Coop *et al.* (2010) proposed and allows to evaluate association of SNP allele frequencies with a population–specific covariable *Z_j_.* Note that *Z_j_* is a (preferably scaled) vector of length *J* containing for each population the measures of interest. Under the STD model, the prior distribution of the vector 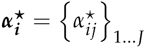 is multivariate Gaussian for each SNP *i*:

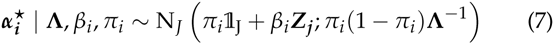

The prior distribution for the correlation coefficients (*β_i_*) is assumed uniform:

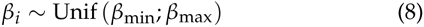

Unless stated otherwise, *β*_min_ = −0.3 and *β*_max_ = 0.3 instead of *β*_min_ = -0.1 and *β*_max_ = 0.1 as in Coop *et al.* (2010).

### The covariate model with auxiliary variable (AUX model)

The AUX model represented in Figure 1C is an extension of the STD model that consists in attaching to each locus regression coefficient *β_i_* a Bayesian (binary) auxiliary variable *δ_i_*. In a similar population genetics context, this modeling was also proposed by Riebler *et al.* (2008) to identify markers subjected to selection in a genome-wide scan for adaptive differentiation (under a 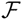–model). In the AUX model, the auxiliary variable actually indicates whether a specific SNP *i* can be regarded as associated with the covariable *Z_j_* (*δ_i_* = 1) or not (*δ_i_* = 0). As a consequence, the posterior mean of *δ_i_* may directly be interpreted as a posterior probability of association of the SNP *i* with the covariable, from which a Bayes Factor (BF) is straightforward to derive (Gautier *et al.* 2009). Under the AUX model, the prior distribution of the vector 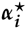 is multivariate Gaussian for each SNP *i*:

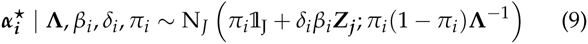

Providing information about marker positions is available, the *δ_i_*’s auxiliary variables also make it easy to introduce spatial dependency among markers. In the context of high–throughput genotyping data, SNPs associated to a given covariable might indeed cluster in the genome due to LD with the underlying (possibly not genotyped) causal polymorphism(s). To learn from such positional information, the prior distribution of *δ* = {*δ_i_*}_(1…_*_I_*_)_, the vector of SNP auxiliary variables, takes the general form of a 1D Ising model with a parametrization inspired from Duforet-Frebourg *et al.* (2014):

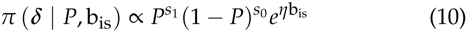
where 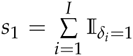 (respectively *s*_0_ = *I* – *s*_1_) are the number of SNPs associated (respectively not associated) with the covariable, and 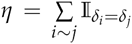 is the number of pairs of consecutive markers (neighbors) that are in the same state at the auxiliary variable (i.e., *δ_i_* = *δ_i_*_+1_). The parameter *P* corresponds to the proportion of SNPs associated to the covariable and is assumed Beta distributed:

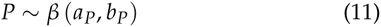

Unless stated otherwise, *a_P_* = 0.02 and *b_P_* = 1.98. This amounts to assume a priori that only a small fraction of the SNPs 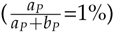 are associated to the covariable, but within a reasonably large range of possible values (e.g., P [*P* > 10%] = 2.8% a priori). Importantly, integrating over the uncertainty on the key parameter *P* allows to deal with multiple testing issues.

Finally, the parameter b_is_, called the inverse temperature in the Ising (and Potts) model literature, determines the level of spatial homogeneity of the auxiliary variables between neighbors. When b_is_ = 0, the relative marker position is ignored (no spatial dependency among markers). This is thus equivalent to assume a Bernoulli prior for the *δ_i_*’s: *δ_i_* ~ Ber(*P*) as in Riebler *et al.* (2008). Conversely, b_is_ > 0 leads to assume that the *δ_i_* with similar values tend to cluster in the genome (the higher the b_is_, the higher the level of spatial homogeneity). In practice, b_is_ = 1 is commonly used and values of b_is_ ≤ 1 are recommended. Note that the overall parametrization of the Ising prior assumes no external field and no weight (as in the so-called compound Ising model) between the neighboring auxiliary variables. In other words, the information about the distances between SNPs is therefore not accounted for and only the relative position of markers are considered. Hence, marker spacing is assumed homogeneous.

## Material and Methods

### MCMC sampler

To explore the different models and estimate the full posterior distribution of the underlying parameters, a Metropolis–Hastings within Gibbs Markov Chain Monte Carlo (MCMC) algorithm was developed (see the File S1 for a detailed description) and implemented in a program called BayPass (for BAYesian Population ASSociation analysis). The software package containing the Fortran 90 source code, a detailed documentation and several example files is freely available for download at http://www1.montpellier.inra.fr/CBGP/software/baypass/. Unless otherwise stated, a MCMC chain first consists in 20 pilot runs of 1,000 iterations each allowing to adjust proposal distributions (for Metropolis and Metropolis–Hastings updates) with targeted acceptance rates lying between 0.2 and 0.4 to achieve good convergence properties (Gilks *et al.* 1996). Then MCMC chains are run for 25,000 iterations after a 5,000 iterations burn-in period. Samples are taken from the chain every 25 post–burn–in iterations to reduce autocorrelations using a so-called thinning procedure. To validate the BayPass sampler, an independent implementation of the core model was coded in the BUGS language and run in the OpenBUGS software (Thomas *et al.* 2009) as detailed in the File S2. Analyses of some (small) test data sets using both implementations gave consistent results (data not shown).

Finally, as a matter of comparison, in the analysis of prior sensitivity in Ω estimation, the BayEnv2 (Günther and Coop 2013) software was also used with default options except the total number of iterations that was set to 50,000.

### Estimation and visualisation of Ω

For BayPass analyses, point estimates of each elements of Ω consisted of their corresponding posterior means computed over the sampled matrices. For BayEnv2 analyses, the first ten sampled matrices were discarded and only the 90 remaining sampled ones were retained. As a matter of comparison, the frequentist estimate of Ω as proposed by Bonhomme *et al.* (2010) and implemented in the FLK package was also considered. Briefly, the FLK relies on a neighbor–joining algorithm on the Reynolds pairwise population distances matrix to build a population tree from which the covariance matrix is deduced (after midpoint rooting of the tree).

For visualization purposes, a given 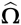 estimate was transformed into a correlation matrix 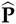 with elements 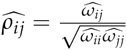 using the `cov2cor()` R function (R Core Team 2015). The graphical display of this correlation matrix was done with the `corrplot()` function from the R package *corrplot* (Wei 2013). In addition, hierarchical clustering of the underlying populations was performed using the `hclust ()` R function considering 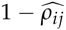 as a dissimilarity measure between each pair of population *i* and *j*. The resulting bifurcating tree was plotted with the `plot. phylo ()` function from the R package *ape* (Paradis *et al.* 2004). Note that the latter representation reduces the correlation matrix into a block-diagonal matrix thus ignoring gene flow and admixture events.

### Computation of the FMD metric to compare Ω matrices

The metric proposed by Förstner and Moonen (2003) for covariance matrices and hereafter referred to as the FMD distance was used to compare the different estimates of Ω and to assess estimation precision and robustness in the prior sensitivity analysis. Let Ω_1_ and Ω_2_ be two (symmetric positive definite) covariance matrices with rank *J*, the FMD distance is defined as:

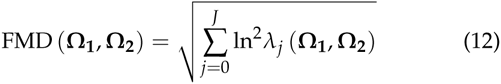
where *λ_j_* (Ω_1_, Ω_2_) represent the *j*th generalized eigenvalue of the matrices Ω_1_ and Ω_2_ that were all computed with the R package *geigen* (Hasselman 2015).

### Computation and calibration of the XtX statistic

Identification of SNPs subjected to adaptive differentiation relied on the XtX differentiation measure introduced by Günther and Coop (2013). This statistic might be viewed as a SNP–specific *F*_ST_ explicitly corrected for the scaled covariance of population allele frequencies. For each SNP *i*, XtX was estimated from the *T* MCMC (post-burn–in and thinned) parameters sampled values, 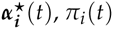 and Λ(*t*), as:

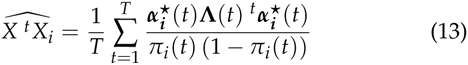

To provide a decision criterion for discriminating between neutral and selected markers, i.e. to identify outlying XtX, we estimated the posterior predictive distribution of this statistic under the null (core) model by analyzing pseudo-observed data sets (POD). PODs are produced by sampling new observations (either allele or read count data) from the core inference model with (hyper–)parameters *a_π_*, *b_π_* and Λ (the most distal nodes in the DAG of Figure 1) fixed to their respective posterior means obtained from the analysis of the original data. The sample characteristics are preserved by sampling randomly (with replacement) SNP vectors of *n_ij_*’s (for allele count data) or *c_ij_*’s (for read count data) among the observed ones. For Pool–Seq data, haploid sample sizes are set to the observed ones. The R (R Core Team 2015) function `simulate .baypass ()` available in the BayPass software package was developed to carry out these simulations. The POD is further analyzed using the same MCMC parameters (number and length of pilot runs, burn-in, chain length, etc.) as for the analysis of the original data set. The XtX values computed for each simulated locus are then combined to obtain an empirical distribution. The quantiles of this empirical distribution are computed and are used to calibrate the XtX observed for each locus in the original data: e.g., the 99% quantile of the XtX distribution from the POD analysis provides a 1% threshold XtX value, which is then used as a decision criterion to discriminate between selection and neutrality. Note that this calibration procedure is similar to the one used in Vitalis *et al.* (2014) for the calibration of their SNP *KLD.*

### Population Association tests and decision rules

Association of SNPs with population–specific covariables is assessed using Bayes Factors (BF) or what may be called “empirical Bayesian P–values” (eBP). Briefly, for a given SNP, BF compares models with and without association while eBP is aimed at measuring to which extent the posterior distribution of the regression coefficient *β_i_* excludes 0. Note that eBP’s are not expected to display the same frequentist properties as classical P–values.

Two different approaches were considered to compute BF’s. The first estimate (hereafter referred to as BF_is_) relies on the Importance Sampling algorithm proposed by Coop *et al.* (2010) and uses MCMC samples obtained under the core model (see File S3 for a detailed description). The second estimate (hereafter referred to as BF_mc_) is obtained from the posterior mean 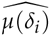 of the auxiliary variable *δ_i_* under the AUX model:

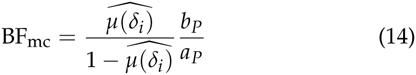
where 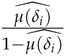 is the (estimated) posterior odds that the locus *i* is associated to the covariable and 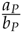 is the corresponding prior odds (Gautier *et al.* 2009). Hereby, BF_mc_ is only derived for the AUX model with b_is_ = 0 (the prior odds being challenging to compute when b_is_ ≠ 0). In practice, to account for the finite MCMC sampled values 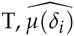 is set equal to 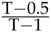 (respectively 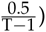) when the posterior mean of the *δ_i_* is equal to 1 (respectively or 0). Note that, through the prior on *P*, the computation of BF_mc_ explicitly accounts for multiple testing issues. BF’s are generally expressed in deciban units (dB) (via the transformation 10log_10_ (BF)). The Jeffreys’ rule (Jeffreys 1961) provide a useful decision criterion to quantify the strength of evidence (here in favor of assocation of the SNP with the covariable) using the following dB unit scale: “strong evidence” when 10 <BF<15, “very strong evidence” when 15<BF<20 and “decisive evidence” when BF>20.

For the computation of eBP’s, the posterior distribution of each SNP was approximated as a Gaussian distribution: 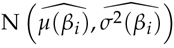 where 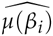 and 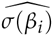 are the estimated posterior mean and standard deviation of the corresponding *β_i_*. The eBP’s are further defined as:

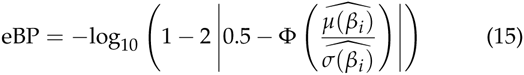
where Φ(*x*) is the cumulative distribution function of the standard normal distribution. Roughly speaking, a value of *β* might be viewed as “significantly” different from 0 at a level of 10^−eBP^%. Two different approaches were considered to estimate the moments of the posterior distribution of the *β_ι_*’s. The first, detailed in the File S3, rely on an Importance Sampling algorithm similar to the one mentioned above and thus uses MCMC samples obtained under the core model. The resulting eBP’s estimates are hereafter referred to as eBP_is_. The second approach relies on posterior samples of the MCMC obtained under the STD model. The resulting eBP’s estimates are hereafter referred to as eBP_mc_.

Note finally, that for estimating BF_mc_ (under the AUX model) and eBP_mc_ (under the STD model), the value of the Λ was fixed to its posterior mean as obtained from an initial analysis carried out under the core model.

### Simulation study

*Simulation under the inference model.* Simulation under the core or the STD inference models defined above (Figure 1) were carried out using the function `simulate .baypass()` available in the BayPass software package. Briefly, a simulated data set is specified by the Ω matrix, the parameters of the Beta distribution for the ancestral allele frequencies (*α_π_* and *b_π_*) and the sample sizes. As a matter of expedience, ancestral allele frequencies below 0.01 (respectively above 0.99) were set equal to 0.01 (respectively 0.99) and markers that were not polymorphic in the resulting simulated data set were discarded from further analyses. For the generation of PODs (see above), the *n_ij_*’s (or the *c_ij_*’s for Pool–Seq data) were sampled (with replacement) from the observed ones and for the power analyses, these were fixed to *n_ij_* = 50 for all the populations. To simulate under the STD model, the simulated *β_i_*’s (SNP regression coefficients) were specified and the population covariable vector Z was simply taken from the standard normal cumulative distribution function such that 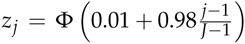 for the *j*th population (out of the *J* ones).

*Individual–based simulations.* Individual–based forward–in–time simulations under more realistic scenarios were carried out under the SimuPOP environment (Peng and Kimmel 2005) as described in de Villemereuil *et al.* (2014). Briefly, three scenarios corresponding to i) an highly structured isolation with migration model (*HsIMM-C*); ii) an isolation with migration model (*IMM*); and iii) a stepping stone scenario (*SS*) were investigated. For each scenario, one data set consisted of 320 individuals belonging to 16 different populations that were genotyped for 5,000 SNPs regularly spread along 10 chromosomes of 1 Morgan length. Polygenic selection acting on an environmental gradient (see de Villemereuil *et al.* 2014, for more details) was included in the simulation model by choosing 50 randomly distributed SNPs (among the 5,000 simulated ones) and affecting them a selection coefficient *s_i_* calculated as a logistic transformation of the corresponding population-specific environmental variable *E_S_* following: 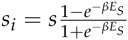 (with *S* = 0.004 and *β* = 5). For each individual, the overall fitness was finally derived from their genotypes using a multiplicative fitness function.

To assess the performance of the AUX model in capturing information from SNP spatial dependency, data sets displaying stronger LD were generated under the *HsIMM-C* (the least favorable scenario, see the Results scetion) by slightly modifying the corresponding script available from de Villemereuil *et al.* (2014). The resulting *HsIMMld-C* data sets each consisted of 5,000 SNPs spread on 5 smaller chromosomes of 4 cM (leading to a SNP density of ca. 1 SNP every 4 kb assuming 1 cM≡1Mb). In the middle of the third chromosome, a locus with strong effect on individual fitness was defined by two consecutive SNPs strongly associated with the environmental covariable (such that *s* = 0.1 and *β* = 1 in the computation of *s_i_,* as defined above). Note that for all the individual–based simulations described in this section, SNPs were assumed in complete Linkage Equilibrium in the first generation.

***comparison with different genome scan methods*** In addition to analyses under the models implemented in BayPass (see above), the *HsIMM-C, IMM* and *SS* individual-based simulated data sets were analyzed with five other popular or recently developed genome scan approaches. These first include BayeScan (Foll and Gaggiotti 2008) which is a Bayesian covariate-free approach that identifies overly differentiated markers (with respect to expectation under a migration–drift equilibrium demographic model) via a logistic regression of the population-by-locus *F*_ST_ on a locus-specific and population-specific effect. The decision criterion was based on a Bayes Factor that quantify the support in favor of a non-null locus effect. Second, the recently developed BayScenv (de Villemereuil and Gaggiotti 2015) model was also used. It is conceived as an extension of BayeScan incorporating environmental information by including a locus-specific regression coefficient parameter (noted *g*) in the above mentioned logistic regression. The decision criterion to assess association with the covariate was based on the estimated posterior probability of *g* being non null. In practice, to limit computation burden for both BayeScan (version 2.1) and BayScenv, default MCMC parameter options of the programs were chosen except for the length of the pilot runs (set to 1,000), the length of the burn-in period (set to 10,000) and the number of sampled values (set 2,500). A third and covariate–free approach consisted in computing the FLK statistics (which might be viewed as the frequentist counterpart of the *X ^t^X* described above) as described in Bonhomme *et al.* (2010). The fourth considered method relied on Latent Factor Mixed Models as implemented in the LFMM (version 1.4) software (Frichot *et al.* 2013) to detect association of allele frequencies differences with population-specific covariables while accounting for population structure via the so-called latent factors. Following de Villemereuil *et al.* (2014) that analyzed the same data sets, the prior number of latent factors required by the program was set to *K* = 15. Note also that *LFMM* analyses were run on individual genotyping data rather than population allele frequencies, which were previously shown to display better performances (de Villemereuil *et al.* 2014). For each data set, the decision criterion to assess association of the SNP with the environmental covariable relied on a P–value that was computed based either on a single analysis (denoted as LFMM) or derived after combining Z–score from 10 independent analyses (denoted as LFMM–10rep) following the procedure described in the LFMM (version 1.4) manual. Finally, the data sets were also analyzed with BayEnv2 (Coop *et al.* 2010) following a two–steps procedure (as required by the program) that was similar to the one performed by de Villemereuil *et al.* (2014). For each data set, a first MCMC of 15,000 iterations was run under default parameter settings and the latest sampled covariance matrix was used as an estimate of Ω. For each SNP in turn, an MCMC of 30,000 iterations was further run to estimate the corresponding *X ^t^ X* and BF based on this latter matrix. To facilitate automation of the whole procedure, a custom shell script was developed.

Each analysis was run on a single node of the same computer cluster to provide a fair comparison of computation times. To further compare the performances of the different models, the actual i) True Positive Rates (TPR) or power i.e. the proportion of true positives among the truly selected loci; ii) False Positive Rates (FPR) i.e. the proportion of false positives among the non selected loci; and iii) False Discovery Rates (FDR) i.e. the proportion of false positives among the significant loci were computed from the analysis of each data set with the different methods for various thresholds covering the range of values of the corresponding decision criterion. From these estimates, both standard Receiver Operating Curves (ROC) plotting TPR against FPR, and Precision-Recall (PR) curves plotting (1-FDR) against TPR could then be drawn.

### Real Data sets

***The HSA_snp_ data set***. This data set is the same as in Coop *et al.* (2010) and was downloaded from the BayEnv2 software webpage (http://gcbias.org/bayenv/). It consists of genotypes at 2,333 SNPs for 927 individuals from 52 human population of the HGDP panel (Conrad *et al.* 2006).

***The BTA_snp_ data set.*** This data set is a subset of the data from Gautier *et al.* (2010b) and consists of 453 individuals from 18 French cattle breeds (from 18 to 46 individuals per breed) genotyped for 42,046 autosomal SNPs displaying an overall MAF>0.01. As detailed in File S4, two breed-specific covariables were considered for association analyses. The first covariable corresponds to a synthetic morphology score (SMS) defined as the (scaled) first principal component of breed average weights and wither heights for both males and females (taken from the French BRG website: http://www.brg.prd.fr/). The second covariable is related to coat color and corresponds to the piebald coloration pattern of the different breeds that was coded as 1 for pied breed (e.g., Holstein breed) and −1 for breeds with a uniform coloration pattern (e.g., Tarine breed).

***The LSA_ps_ data set***. This data set was obtained from whole transcriptomes of pooled Littorina saxatilis (LSA) individuals belonging to 12 different populations originating (Westram *et al.* 2014). These populations originate from three distinct geographical regions (UK, the United Kingdom; SP, Spain and SW, Sweden) and lived in two different ecotypes corresponding to the so-called “wave” habitat (subjected to wave action) and “crab” habitat (i.e., subjected to crab predation). The *mpileup* file with the aligned RNA–seq reads from the 12 pools (three countries × two ecotypes × two replicates) onto the draft LSA genome assembly was downloaded from the Dryad Digital Repository `doi: 10.5061/dryad. 21pf0` (Westram *et al.* 2014). The *mpileup* file was further processed using a custom awk script to perform SNP calling and derive read counts for each alternative base (after discarding bases with a BAQ quality score <25). A position was considered as variable if i) it had a coverage of more than 20 and less than 250 reads in each population; ii) only two different bases were observed across all the five pools and; iii) the minor allele was represented by at least one read in two different pool samples. Note that tri–allelic positions for which the two most frequent alleles satisfied the above criteria and with the third allele represented by only one read were included in the analysis as bi-allelic SNPs (after filtering the third allele as a sequencing error). The final data set then consisted of allele counts for 53,387 SNPs. As a matter of expedience, the haploid sample size was set to 100 for all the populations because samples consisted of pools of ca. 40 females with their embryos (from tens to hundreds per female) (Westram *et al.* 2014). To carry out the population analysis of association with ecotype and identify loci subjected to parallel phenotypic divergence, the habitat is considered as a binary covariable respectively coded as 1 for the “wave” habitat and −1 for the “crab” habitat.

## Results

### Performance of the core model for estimation of the scaled population covariance matrix Ω

The scaled covariance matrix Ω of population allele frequencies represents the key parameter of the models considered in this study. Evaluating the precision of its estimation is thus crucial. To illustrate how prior parametrization might influence estimation of Ω, we first analyzed the BTA_snp_ (with J=18 French cattle populations) and the HSA_snp_ (with J=52 worldwide human populations) data sets using both BayPass (under the core model represented in Figure 1A with *ρ* = 1) and BayEnv2 (in which *ρ* = *J* and *a_π_* = *b_π_* = 1 according to Coop *et al.* (2010)). Note that the sampled populations in these two data sets have similar characteristics in terms of the overall *F_ST_* (*F_ST_* = 9.84% and *F_ST_* = 10.8% for the cattle and human sampled populations respectively). The resulting estimated Ω matrices are hereafter denoted as 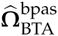 and 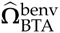 respectively for the cattle data set and are represented in Figure 2. Similarly, for the human data set, the resulting 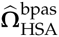 and 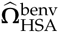 are represented in Figure S1. For both data sets, the comparisons of the two different estimates of Ω reveal clear differences that suggest in turn some sensitivity of the model to the prior assumption. Analyses under three other alternative BayPass model parameterizations (i) *ρ* = 1 and *a_π_* = *b_π_* = 1; ii) *ρ* = *J* and; iii) *ρ* = *J* and *a_π_* = *b_π_* = 1) confirmed this intuition (Figure S2). For the human data set, the FMD between the different estimates of Ω varied from 1.73 (BayPass with *ρ* = 1 *vs* BayPass with *ρ* = 1 and *a_π_* = *b_π_* = 1) to 31.1 (BayPass with *ρ* = 1 *vs* BayPass with *ρ* = 52). However, for the cattle data set that contains about 20 times as many SNPs for 3 times less populations, the four BayPass analyses gave consistent estimates (pairwise FMD always below 0.5) that clearly depart from the BayEnv2 one (pairwise FMD always above 14). Note also that BayPass estimates were in better agreement with the historical and geographic origins of the sampled breeds (see Figure 2 and Gautier *et al.* (2010b) for further details).

**Figure 2.**
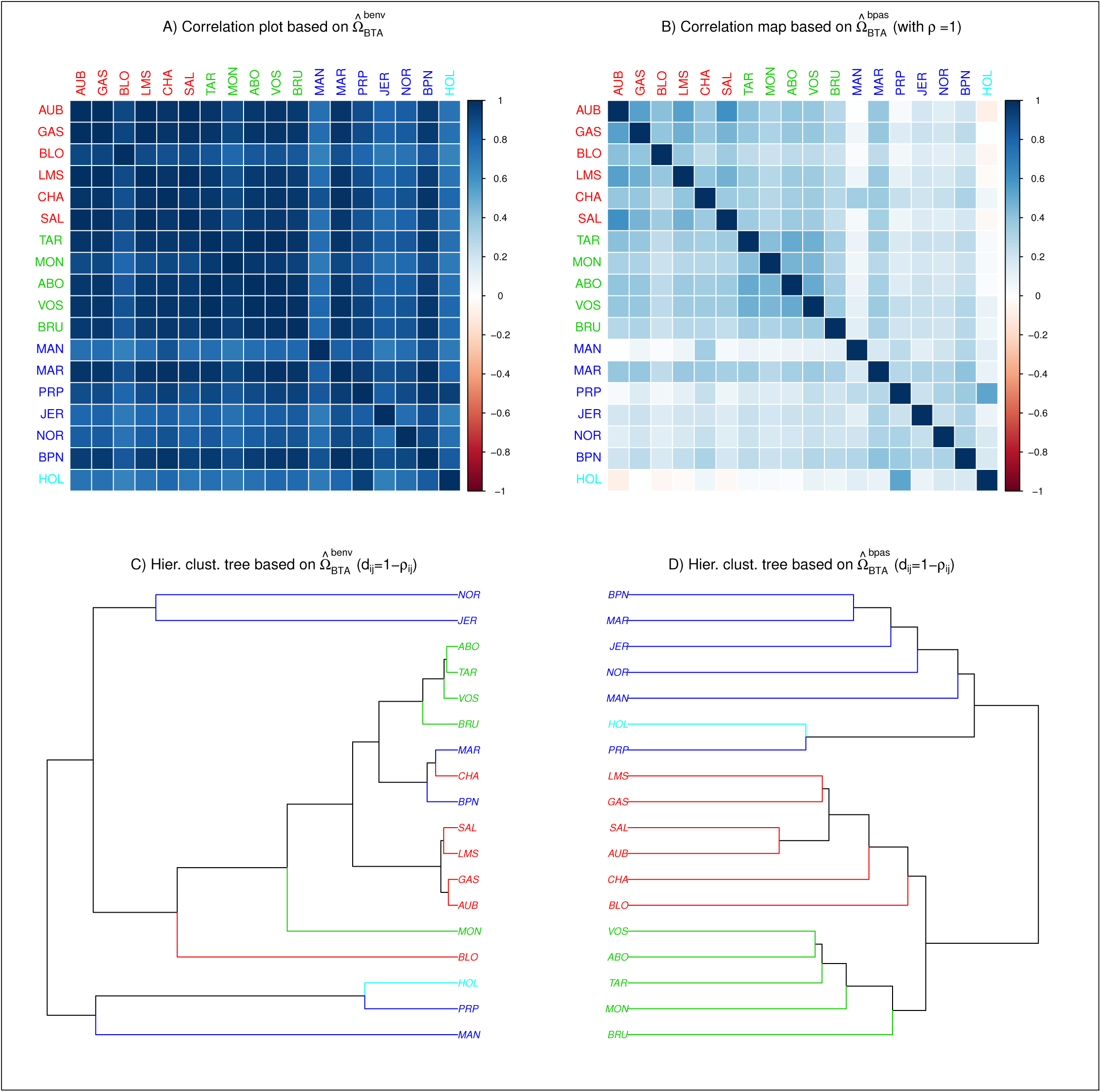
Representation of the scaled covariance matrices Ω among 18 French cattle breeds 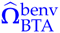 (A and C) as estimated from BayEnv2 (Coop *et al.* 2010) and 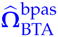 (B and D) as estimated from BayPass under the core model with *ρ* = 1. Both estimates are based on the analysis of the BTA_snp_ data set consisting of 42,036 autosomal SNPs (see the main text). Breed codes (and branches) are colored according to their broad geographic origins (see File S4 and Gautier *et al.* (2010b) for further details) with populations in red, blue and green originating from South-Western and Central France, North-Western France, and Eastern France (e.g. Alps).

Overall these contrasting results call for a detailed analysis of the sensitivity of the model to prior specifications on both Ω (*ρ* value) and the *π_i_* Beta distribution parameters (*a_π_* and *b_π_*), but also to data complexity (number and heterozygosity of SNPs). To that end we first simulated under the core inference model (Figure 1A) data sets for four different scenarios labeled SpsH1, SpsH2, SpsB1 and SpsB2. In SpsH1 and SpsH2 (respectively SpsB1 and SpsB2), the population covariance matrix was set equal to 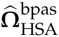 (respectively 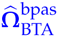), and in SpsH1 and SpsB1 (respectively SpsH2 and SpsB2) the *π_i_*’s were sampled from a Uniform distribution over (0,1) (respectively a *Beta* (0.2, 0.2) distribution). Note that the two different *π_i_* distributions lead to quite different SNP frequency spectrum, the Uniform one approaching (ascertained) SNP chip data (i.e., good representation of SNPs with an overall intermediate MAF) while the *Beta* (0.2, 0.2) one is more similar to that obtained in whole genome sequencing experiments with an over–representation of poorly informative SNPs (see, e.g., results obtained on the LSA_ps_ Pool–Seq data below). To assess the influence of the number of genotyped SNPs, data sets consisting of 1,000, 5,000, 10,000 and 25,000 SNPs were simulated for each scenario. For each set of simulation parameters, ten independent replicate data sets were generated leading to a total of 160 simulated data sets (10 replicates × 4 scenarios × 4 SNP numbers) that were each analyzed with BayEnv2 (Coop *et al.* 2010) and four alternative BayPass model parameterizations (i) *ρ* = 1; ii) *ρ* = 1 and *a_π_* = *b_π_* = 1; iii) *ρ* = *J* and; iv) *ρ* = *J* and *a_π_* = *b_π_* = 1). As a matter of comparisons, the FLK frequentist estimate (Bonhomme *et al.* 2010) of the covariance matrices was also computed. FMD distances (averaged across replicates) of the resulting Ω estimates from their corresponding true matrices are represented in Figure 3. Note that for a given simulation parameter set, the FMD distances remained quite consistent (under a given model parametrization) across the ten replicates (Figure S3).

**Figure 3.**
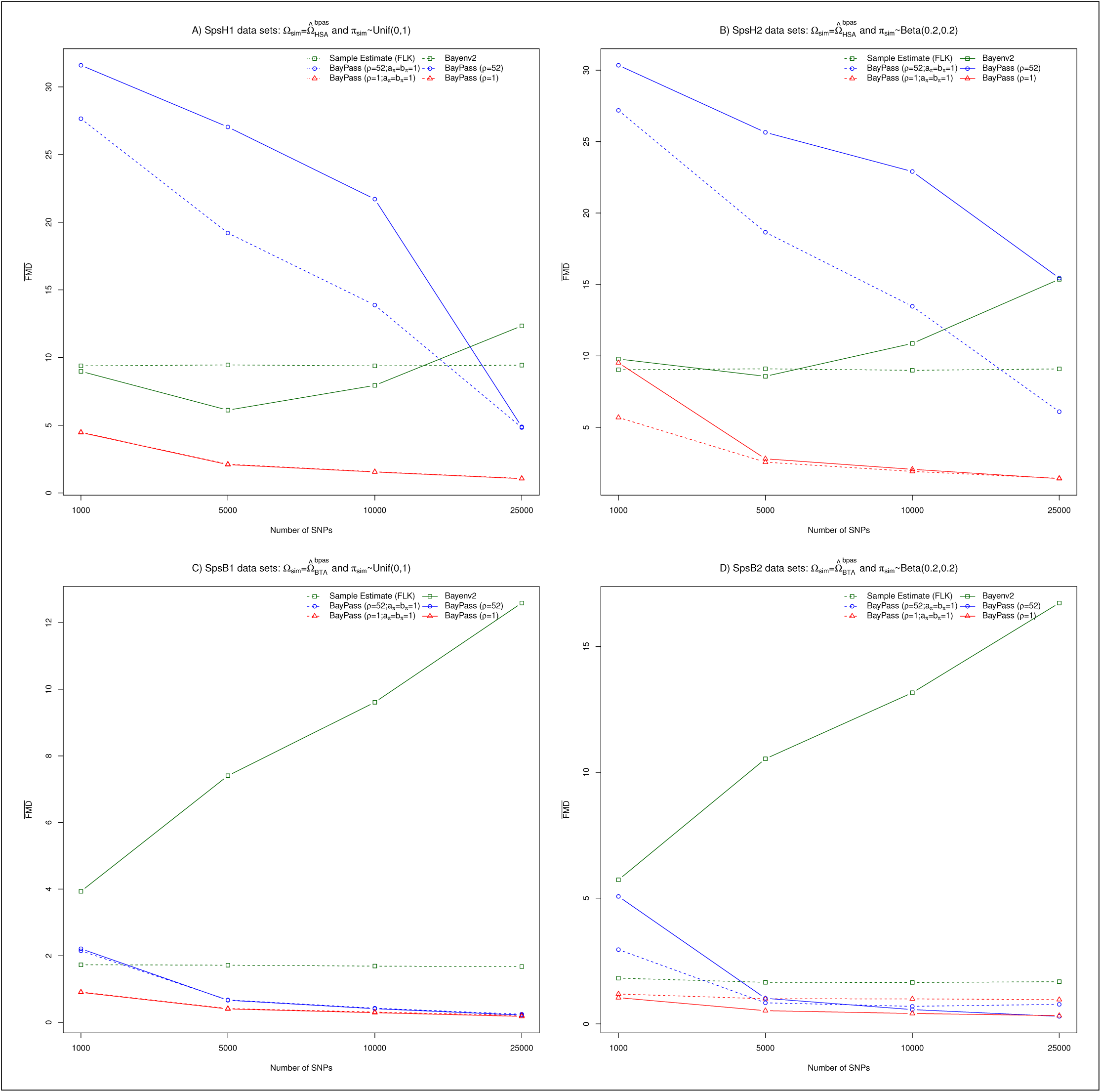
FMD distances (Förstner and Moonen 2003) between the matrices used to simulate the data sets and their estimates. Simulation scenarios are defined according to the matrix Ω_sim_ used to simulated the data (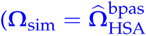 in A and B; and 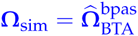 in C and D) and the sampling distribution of the *π_i_*’s (Unif(0,1) in A and C and Beta(0.2,0.2) in B and D). For each scenario, ten independent data sets of 1,000, 5,000, 10,000 and 25,000 markers were simulated (160 data sets in total) and analyzed with B**ay**Env2 (Coop *et al.* 2010) and four alternative BayPass model parameterizations (i) *ρ* = 1; ii) *ρ* = 1 and *a_π_ = b_π_* = 1; iii) *ρ = J* and ; iv) *ρ = J* and *a_π_ = b_n_* = 1). As a matter of comparisons, the FLK frequentist estimate(Bonhomme *et al.* 2010) of the covariance matrices was also computed. Each point in the curves is the average of the ten pairwise FMD distances between the underlying Ω_sim_ and each of the 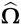 estimated in the ten corresponding simulation replicates.

Except for the BayEnv2 and FLK analyses, the estimated matrices converged to the true ones as the number of SNPs (and thus the information) increase. In addition, as observed above for real data sets, the BayEnv2 estimates were always quite different from those obtained with BayPass parametrized under the same model assumptions (*ρ* =npop and *a_π_* = *b_π_* = 1). It should also be noticed that reproducing the same simulation study by using the 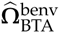 and 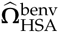 matrices in the four different scenarios lead to similar patterns (Figure S4). Reasons for this behavior of BayEnv2 (possibly the result of some minor implementation issues) were not investigated further and we hereafter only concentrated on results obtained with BayPass.

As expected, the optimal number of SNPs also depends on their heterozygosity. Hence, when the simulated *π_i_*’s were sampled from a *Beta* (0.2, 0.2) (Figure 3B and D) instead of a Unif(0,1) distribution, a higher number of SNPs was required (compare Figures 3B and A; and Figures 3D and C, respectively) to achieve the same accuracy. Likewise, all else being equal, the estimation precision was found always lower for the SpsH1 (and SpsH2) than SpsB1 (and SpsB2) scenarios. This shows that the optimal number of SNPs is an increasing function of the number of sampled populations. One might also expect that more SNPs are required when population differentiation is lower (although this was not formally tested here). Regarding the sensitivity of the models to the prior definition, the parametrization with *ρ* = 1 clearly outperformed the more informative one (*ρ* = *J*), most particularly for smaller number of SNPs and more complex data sets. Naturally, estimating the parameters *a_π_* and *b_π_* compared to setting them to *a_π_* = *b_π_* = 1 had almost no effect in the estimation precision of Ω for the SpsH1 and SpsB1 scenarios, their resulting posterior means being slightly larger than one (≃ 1.1 due probably to the simulation SNP ascertainment scheme as described in Material and Methods). Interestingly however, a substantial gain in precision was obtained for the SpsH2 and SpsB2 data sets (for which 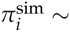 ~ *Beta* (0.2, 0.2)). Hence, for the SpsB2 data sets (Figure 3D), the FMD curves reached a plateau with the *a_π_ = b_π_* = 1 parametrization (for both *ρ* = 1 and *ρ* = 18) as the number of SNPs increase whereas precision kept improving when *a_π_* and *b_π_* were estimated.

We finally investigated to which extent estimation of *a_π_* and *b_π_* might improve robustness to SNP ascertainment. To that end, ten additional independent data sets of 100,000 SNPs were simulated under both the SspH1 and SspB1 scenarios. For each of the twenty resulting data sets, six subsamples were constituted by randomly sampling 25,000 SNPs with an overall MAF>0, >0.01, >0.025, >0.05, >0.075 and >0.10 respectively. The 120 resulting data sets (2 scenarios × 10 replicates × 6 MAF thresholds) were analyzed with BayPass (assuming *ρ* = 1) by either estimating *a_π_* and *b_π_* or setting *a_π_ = b_π_* = 1. Although the estimation precision of Ω was found to decrease with increasing MAF thresholds (Figure S5), estimating *a_π_* and *b_π_* allowed to clearly improve accuracy in these examples. Note however, that the effect of the ascertainment scheme remained limited, in particular for small MAF thresholds (MAF<0.05).

### Performance of the XtX statistics to detect overly differentiated SNPs

To evaluate the performance of the XtX statistics to identify SNPs subjected to selection, data sets were simulated under the STD inference model (Figure 1B), i.e., with a population–specific covariable. This simulation strategy was mainly adopted to compare covariable-free XtX based decision (scan for differentiation) with association analyses (based on covariate models) as described in the next section. Obviously, the XtX is a covariable-free statistic that is powerful to identify SNPs subjected to a broader kind of adaptive constraints, as elsewhere demonstrated (Bonhomme *et al.* 2010; Günther and Coop 2013). Hence, two different (demographic) scenarios, labeled SpaH and SpaB, were considered. In the scenario SpaH (respectively SpaB), Ω^sim^ was set equal to 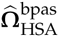 (respectively 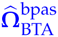), and the *π_i_*’s were sampled from a Uniform distribution. For each scenario, 25,600 SNPs were simulated of which 25,000 are neutral SNPs (i.e., with a regression coefficient *β_i_* = 0) and 600 are SNPs associated with a normally distributed population–specific covariable (see Material and Methods) and with regression coefficients *β_i_* = −0.2 (n=100), *β_i_* = −0.1 (n=100), *β_i_* = −0.05 (n=100), *β_i_* = 0.05 (n=100), *β_i_* = 0.1 (n=100),*β_i_* = 0.2 (n=100). For each scenario, ten independent replicate data sets, each with a randomized population covariable vector, were generated. The resulting 20 simulated data sets (10 replicates × 2 scenarios) were then analyzed with four alternative BayPass model parameterizations corresponding to i) the core model (Figure 1A) with *ρ* = 1; ii) the core model by setting Ω = Ω^sim^; iii) the STD model (Figure 1B) by setting Ω = Ω^sim^ and; iv) the default AUX model (Figure 1C) i.e. with b_is_ = 0 and Ω = Ω^sim^.

As expected, under the core model, the higher |*β*_i_| the higher the estimated XtX on average (Figure S6). As a matter of expedience, for power comparisons, 1% POD threshold were further defined for each analysis using the XtX distribution obtained for SNPs with simulated *β_i_* = 0. Note that the resulting thresholds were very similar to those obtained using independent data sets (e.g., SpsH1 and SpsB1) that lead to FPR close to 1%. As shown in Table 1, the power was optimal (> 99.9%) for strongly associated SNPs (|*β_i_*| = 0.2) in both scenarios but remained small (< 10%) for weakly associated SNPs. In addition, power was always higher with the SpaH than with the SpaB data probably due to a more informative design (three times as many populations). Likewise, estimating Ω (i.e., including information from the associated SNPs) slightly affected the performance of the XtX-based criterion when compared to setting Ω = Ω^sim^ (see Table 1 and also the ROC curve analyses in Figure S7). Yet the resulting estimated matrices Ω were close to the true simulated ones (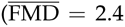 across the SpaH and 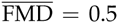 across the SpaB simulated data sets) suggesting in turn that the core model is also robust to the presence of SNPs under selection (at least in moderate proportion). Conversely, a misspecification of the prior Ω, as investigated here by similarly analyzing the SpaH (respectively SpaB) data sets under the core, the STD and the AUX models but setting 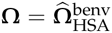 (respectively 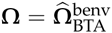), lead to an inflation of the XtX estimates (Figure S8). The XtX mean was in particular shifted away from *J* (number of populations) expected under neutrality (see also Figure 5 in Günther and Coop (2013)). As a consequence, the overall performances of the XtX-based criterion were clearly impacted (see Table S1 and ROC in Figure S7).

Interestingly, under both the STD and AUX models, the distribution of the XtX for SNPs associated to the population covariable was similar to the neutral SNP one, whatever the underlying *β_i_* (Figure S6). Accordingly, the corresponding true positive rates were close to the nominal POD threshold in Table 1. This suggests that both covariate models allow to efficiently correct the XtX estimates for the (“fixed”) covariable effect of the associated SNPs.

**Table 1.**
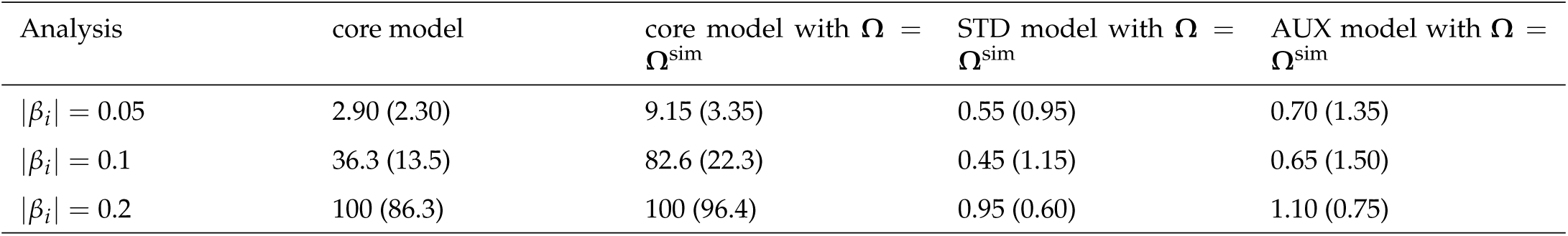
True Positive Rates (TPR) at the 1% POD threshold as a function of the simulated |*β_i_*| values for four different model parameterizations. TPR are given in % and were computed by combining results over the ten replicate data sets for each SpaH (and SpaB given in parenthesis) scenarios.

| Analysis | core model | core model with $\Omega = \Omega^{\text{sim}}$ | STD model with $\Omega = \Omega^{\text{sim}}$ | AUX model with $\Omega = \Omega^{\text{sim}}$ |
| --- | --- | --- | --- | --- |
| $\vert \beta_i\vert = 0.05$ | 2.90 (2.30) | 9.15 (3.35) | 0.55 (0.95) | 0.70 (1.35) |
| $\vert \beta_i\vert = 0.1$ | 36.3 (13.5) | 82.6 (22.3) | 0.45 (1.15) | 0.65 (1.50) |
| $\vert \beta_i\vert = 0.2$ | 100 (86.3) | 100 (96.4) | 0.95 (0.60) | 1.10 (0.75) |

### Performance of the models to detect SNP associated to a population–specific covariable

The performances of the STD and AUX models to identify SNPs associated to a population–specific covariable were further evaluated using results obtained on the SpaH and SpaB data sets (see above). As shown in Figure 4, the Importance Sampling estimates of the *β_i_* coefficients (computed from parameter values sampled under the core model) were found less accurate than posterior mean estimates obtained from values sampled under the STD or AUX models. For smaller |*βi*| however, the introduction of the auxiliary variable (AUX model) tended to shrink the estimates towards zero in the SpaB data sets probably due, here also, to a less powerful design (three times less populations).

**Figure 4.**
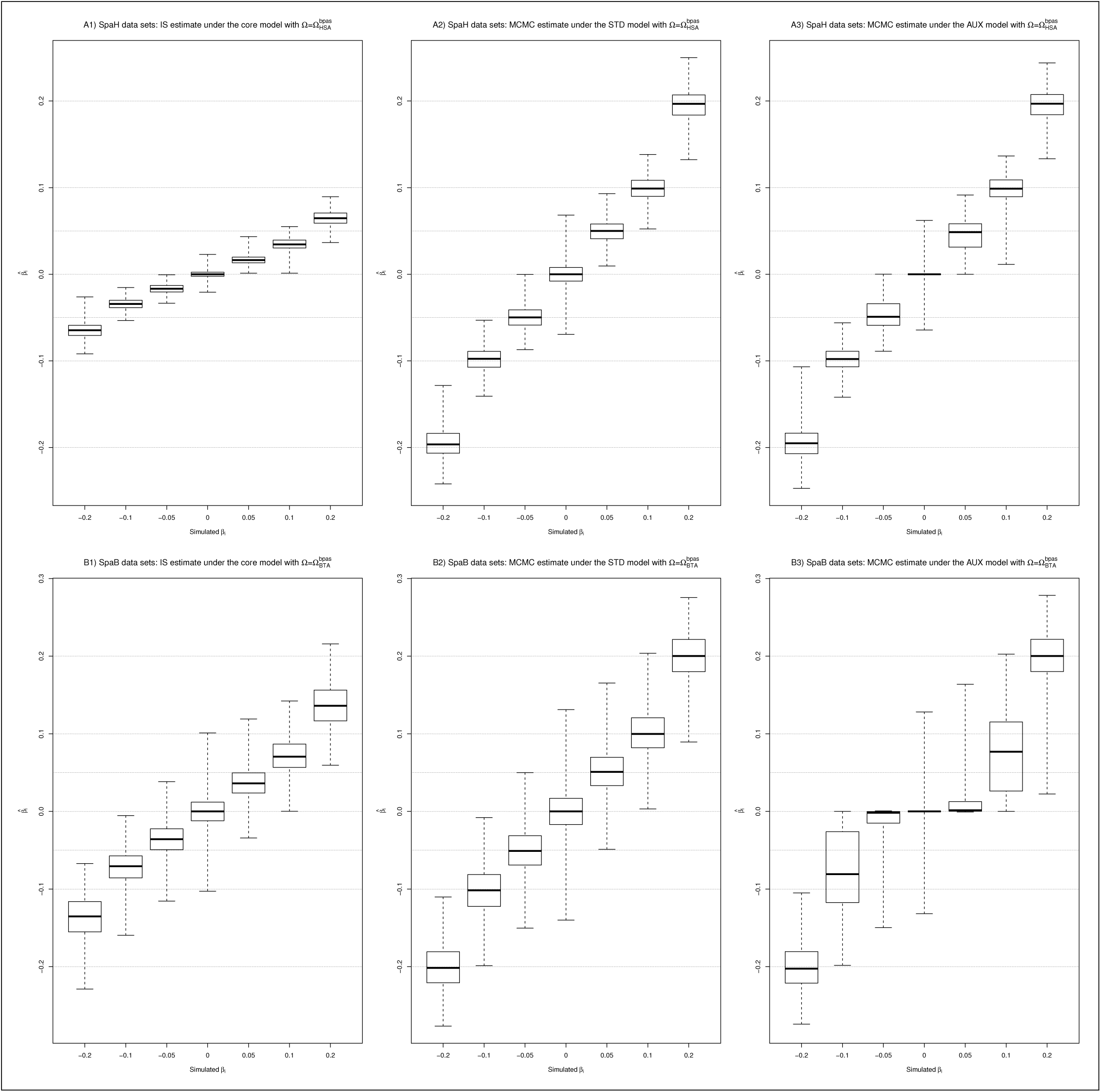
Distribution of the estimated SNP regression coefficients *β_i_* as a function of their simulated values obtained from analyses under three different model parameterizations with 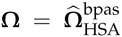 (for SpaH data) and 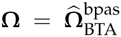 (for SpaB data). For a given scenario (SpaH and SpaB), results from the ten replicates are combined.

Accordingly, the BF’s estimated under the AUX model (BF_mc_) had more power to identify SNPs associated to the population-specific covariables than the corresponding BF_is_ (Table 2 and Figure S9). Indeed, although constrained by construction to a maximal value (here 53.0 dB) that both depends on the number of MCMC samples (here 1,000) and on the prior expectation of *P* (here 0.01), at the “decisive evidence” threshold of 20 dB (Jeffreys 1961), the TPR for SNPs with a simulated |*β_i_*| = 0.05 were for instance equal to 81.7% with BF_mc_ for the SpaH data compared to 31.9% with the BF_is_ based decision criterion (Table 2). For the SpaH data (but not for the SpaB ones) a similar trend was observed when comparing decision criteria based on the eBP_is_ (relying on Importance Sampling algorithm) and the eBP_mc_ as estimated under the STD model (see Table 2 and Figure S10). In addition, Table 2 shows that the intuitive, but still arbitrary, threshold of 3 on the eBP performed worse than the 20 dB threshold on the BF, particularly for the smallest |*β_i_*|. This suggests that a decision criterion rule relying on the BFmc may be the most reliable in the context of these models.

**Table 2.**
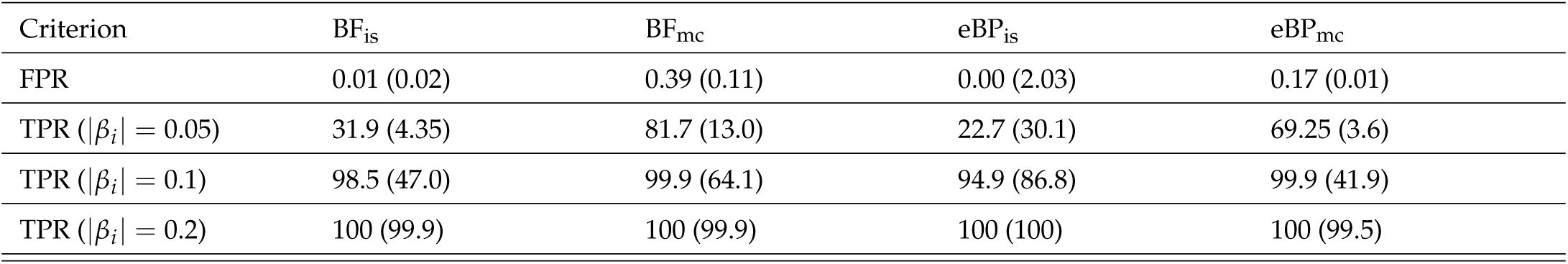
True (TPR) and False (FPR) Positive Rates as a function of the decision criterion and the model parametrization (with 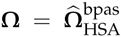 for the SpaH and 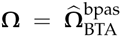 for the SpaB data sets respectively). The thresholds are set to 20 dB for both the BF_is_ and BF_mc_ Bayes Factors; and to 3 for both the eBP_is_ and eBP_mc_ (empirical) Bayesian P-values. The true and false positive rates (given in %) are computed by combining results over the ten replicate data sets from the SpaH and SpaB (given in parenthesis) scenarios.

We next explored how a misspecification of the prior Ω affected the estimation of the *β_i_*’s and the different decision criteria. As in the previous section, we considered results obtained for the SpaH (respectively SpaB) data sets with analyses setting 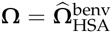 (respectively 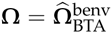). Surprisingly, although the Importance Sampling estimates of the *β_i_*’s obtained under the core model clearly performed poorer (particularly for the SpaB data), the estimates obtained under the STD and AUX models were not so affected (Figure S11). Nevertheless, if the resulting TPR and FPR were similar to the previous ones for the SpaH data, for the SpaB data the power to detect associated SNPs strongly decreased with both the BF_is_ and eBP_is_ criteria. Conversely, increased FPR were observed with the BF_mc_ (up to 22.5%) and eBP_mc_ based decision criteria (see Table S2 and compare with Table 2). These results thus suggest that the influence of model misspecification, although unpredictable, may be critical for association studies under the STD and AUX covariate models.

### Comparison of the performances of BayPass with other genome–scan methods under realistic scenarios

To compare the performances of the different approaches implemented in BayPass with other popular or recently developed methods, data sets simulated under three realistic scenarios were considered. Following de Villemereuil *et al.* (2014) (see Material and Methods), these correspond to i) an highly structured isolation with migration model (*HsIMM–C*); ii) an isolation with migration model (*IMM*); and iii) a stepping stone scenario (*SS*) with polygenic selection acting on an environmental gradient. In total 300 data sets (100 per scenario), each consisting of genotyping data on 5,000 SNPs for 320 individuals belonging to 16 different populations were analyzed with BayPass under the core model (to estimate XtX, BF_is_ and eBP_is_), the STD model (to estimate eBP_mc_ and also the XtX corrected for the fixed covariable effect) and the AUX model (to estimate BF_is_ and also the corrected XtX). These data sets were also analyzed with five other programs (see Material and Methods), two of which, namely BayeScan (Foll and Gaggiotti 2008) and FLK (Bonhomme *et al.* 2010), implementing (only) covariate–free approaches; and the three others, namely BayEnv2 (Coop *et al.* 2010), LFMM (Frichot *et al.* 2013) and BayScenv (de Villemereuil and Gaggiotti 2015) allowing to test for association with population–specific covariable.

For each scenario, average ROC and PR curves resulting from the analyses of the 100 simulated data sets are plotted for the different methods (and decision criteria) in Figure 5. In addition, Area Under the ROC Curve (AUC) together with averaged computation times are detailed in Table 3. In agreement with previous studies (e.g., de Villemereuil *et al.* 2014), under such complex scenarios with polygenic selection, the association based methods clearly outperformed covariable–free approaches (BayeScan, FLK and XtX–based criterion). For the latter however, the BayPass XtX (as estimated under the core model) always performed better than BayeScan and FLK in all the three scenarios. Surprisingly, for the *HsIMM–C* and the *SS* scenarios, the BayEnv2 XtX–based criterion lead to higher AUC than its BayPass counterpart with a value close to the BayEnv2 BF association test (Table 3).

**Figure 5.**
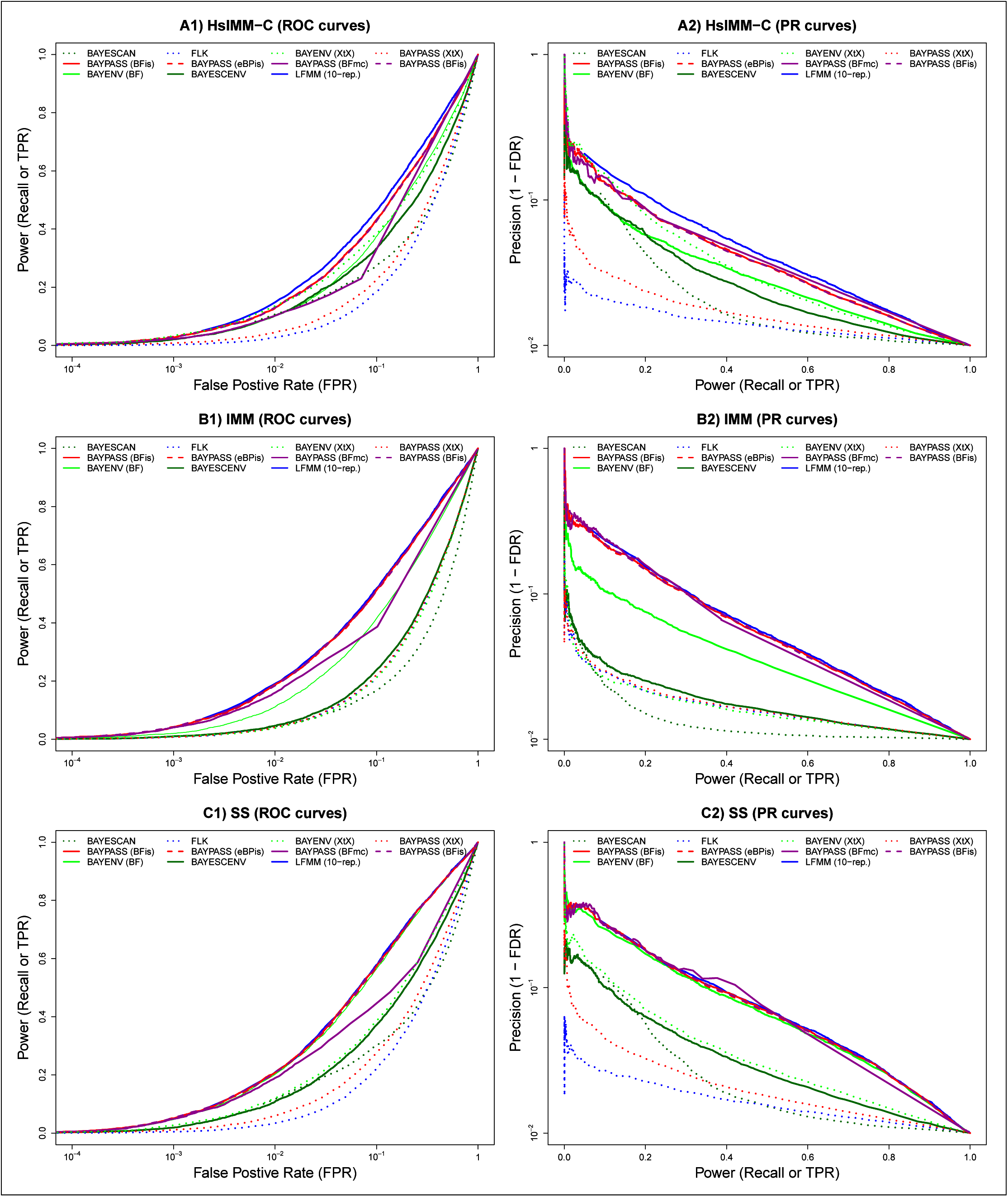
Comparison of the performances of BayPass with other genome-scan methods based on data simulated under three different scenarios (*HsIMM–C, IMM* and *SS*) that include polygenic selection. For each scenario, ROC and PR curves corresponding to the different approaches (and decision criteria) were plotted from the actual TPR, FPR and FDR estimates averaged over the results of 100 independent data sets.

Among the association–based methods, BayPass was found to display similar performances (using the BF_is_, eBP_is_ and eBP_mc_) than *LFMM*–10rep, being even slightly better than single-run *LFMM* analyses for the *IMM* and *SS* scenarios. Both methods outperformed BayeScenv and BayEnv2 in all scenarios (except *SS* scenario for the latter). It should be noted that *LFMM*–10rep analyses were based on individual genotyping data (and a balanced design) which represent the most favorable situation (de Villemereuil *et al.* 2014). The BF_mc_ criterion displayed similar performances in the PR analysis than the BayPass BF_is_, eBP_is_ and eBP_mc_ criteria. Nevertheless, ROC AUC values were always found lower when considering BF_mc_ probably as a result of the inherent correction in the AUX model for multiple testing issues which, as expected, affects the power. Interestingly, as expected from previous results, the XtX calculated under the STD model (and to a lesser extent the AUX model) lead here to a worthless decision criterion (ROC AUC almost equal to 0.5) illustrating the efficiency of the correction for the fixed covariable effect (Table 3).

**Table 3.** Computation times and Area Under the ROC Curves (AUC in %) for the analyses of the *HsIMM-C, IMM* and *SS* data sets using the different genome–scan approaches. Computation times are averaged over the 300 analyses (100 data sets times 3 scenarios)

| Method | Criterion | Mean (median)<br>Computation<br>Time in min | *HsIMM-C* | *IMM* | *SS* |
| --- | --- | --- | --- | --- | --- |
| BAYESCAN | BF | 529 (469) | 60.13 | 53.81 | 62.05 |
| FLK | FLK | 0.16 (0.16) | 58.92 | 61.63 | 62.17 |
| BayEnv2 | XtX | 660 (358) | 70.45 | 61.00 | 72.16 |
|  | BF |  | 70.58 | 73.84 | 81.96 |
| BAYPASS (core model) | XtX | 22.6 (22.2) | 61.66 | 61.88 | 65.33 |
|  | Bfis |  | 74.36 | 78.91 | 82.29 |
|  | eBPis |  | 74.33 | 78.78 | 82.22 |
| BAYPASS (STD model) | XtX | 21.4 (17.8)<sup>a</sup> | 49.85 | 49.16 | 47.72 |
|  | eBPmc |  | 74.15 | 78.76 | 82.22 |
| BAYPASS (AUX model) | XtX | 45.3 (44.9)<sup>a</sup> | 60.60 | 59.82 | 61.08 |
|  | Bfmc |  | 58.30 | 65.24 | 70.51 |
| BAYESCENV | Post. Prob. | 510 (478) | 66.93 | 62.34 | 70.36 |
| LFMM<sup>b</sup> | P-value | 33.0 (30.4)<sup>c</sup> | 75.58 | 78.29 | 81.98 |
| LFMM-10rep<sup>b</sup> | P-value | 310 (248)<sup>c</sup> | 76.27 | 79.37 | 82.56 |

<sup>a</sup> not accounting for the time required to estimate the covariance matrix (obtained here after running BAYPASS under the core model)
<sup>b</sup> Analyses were carried out using individual genotyping data rather than (population) allele count which provides the best performances (see, e.g., de Villemereuil et al. 2014)
<sup>c</sup> Not accounting for the time required to estimate the number of latent factor K (set here to K=15)

Finally, under the parameter options chosen to run the different programs (see, Material and Methods), BayPass analyses were always among the most computational efficient approaches (Table 3). For instance, under the core model, BayPass was found to run 1.5 times faster than a single *LFMM* run.

### Performance of the Ising prior to account for SNP spatial dependency in association analyses

To evaluate the ability of the AUX model Ising prior to capture SNP spatial dependency information, 100 data sets simulated under the *HsIMMld-C* scenario (see Material and Methods) were analyzed under the AUX model with three different parameterizations for the Ising prior i) b_is_ = 0 (no spatial dependency); ii) b_is_ = 0.5 and; iii) b_is_ = 1. For each data set, analyses with and without the causal variants were carried out and the required estimate of the covariance matrix was obtained from a preliminary analysis performed under the core model. As shown in Figure 6, increasing b_is_ improved the mapping precision. Indeed, both a noise reduction at neutral position and a sharpening of the 95% envelope (containing 95% of the *δ_i_* posterior means across the 100 simulated data sets) around the selected locus can be observed (e.g., compare Figure 6 A1 and A3). Interestingly, given the considered SNP density (and level of LD) excluding the causal variants had only marginal effect on the overall results.

**Figure 6.**
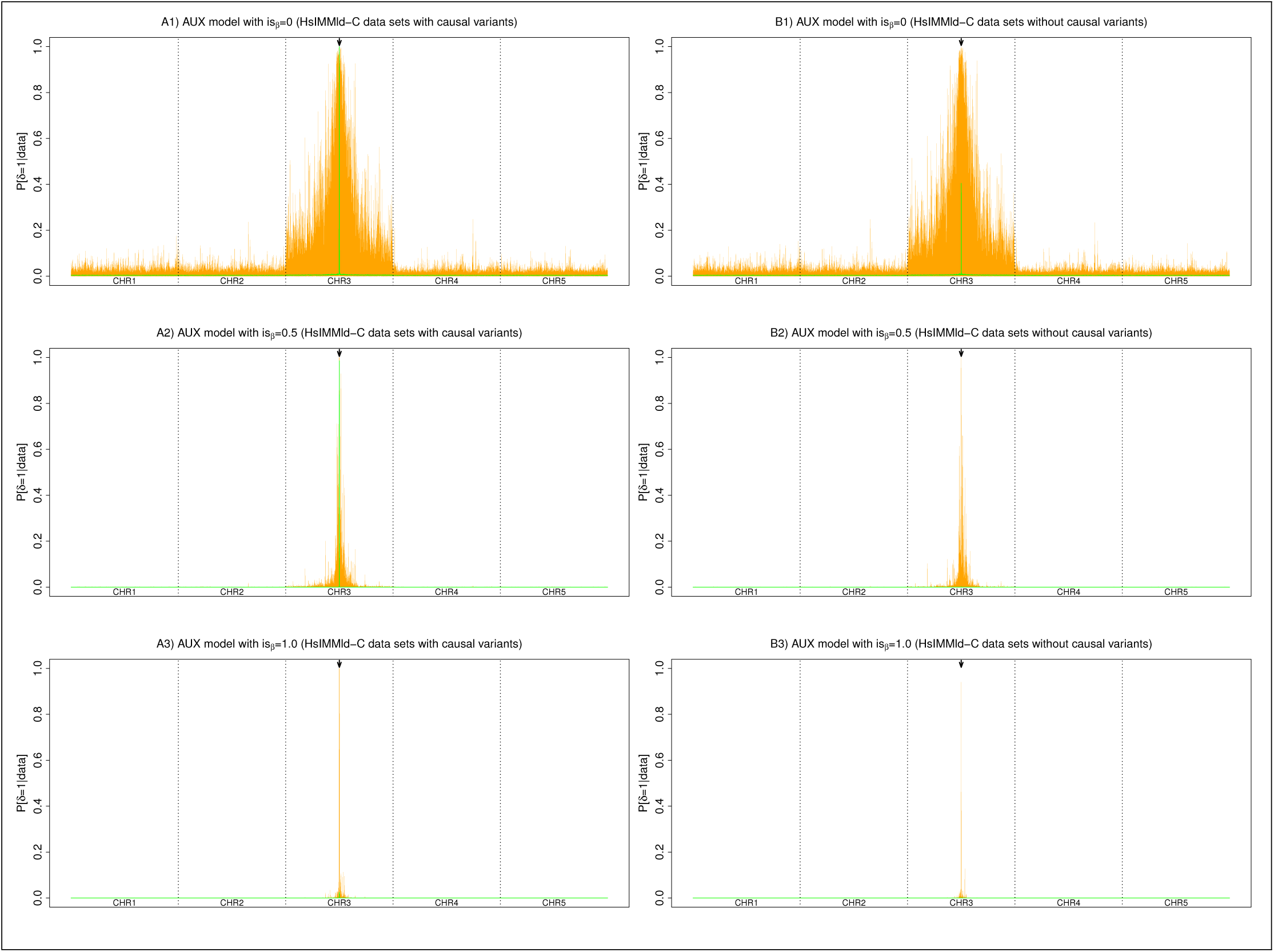
Comparison of the performances of three different Ising prior parameterizations for the AUX model (b_is_ = 0, b_is_ = 0.5 and b_is_ = 1) on the *HsIMMld-C* simulated data sets with (A1, A2 and A3) and without (B1, B2 and B3) the causal variants. Each panel summarizes the distribution at each SNP position (x-axis) of the *δ_i_* (auxiliary variable) posterior means over the 100 independent simulated data with the median values in green and the 95% envelope in orange. Each simulated data set consisted of 5,000 SNPs spread on 5 chromosome of 4 cM. In the middle of the third chromosome (indicated by an arrow), a locus with a strong effect on individual fitness was defined by two consecutive SNPs strongly associated with the environmental covariable.

### Analysis of the French cattle SNP data

The XtX estimates were obtained for the 42,046 SNPs of the *BTA*_snp_ data (Figure S12) from the previous analysis under the core model with *ρ* = 1 (e.g., Figure 2). In agreement with above results, setting instead 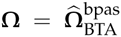 (the estimate of Ω obtained in the latter analysis) gave almost identical XtX estimates (*r* = 0.995). To calibrate the XtX’s, a POD containing 100,000 simulated SNPs was generated and further analyzed leading to a posterior estimate of Ω very close to 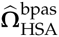(*FMD* = 0.098). Similarly, the posterior means of *a_π_* and *b_π_* obtained on the POD data set (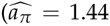 and 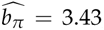, respectively) were almost equal to the ones obtained in the original analysis of the *BTA*_snp_ data set (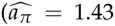 and 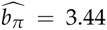, respectively). This indicated that the POD faithfully mimics the real data set, allowing the definition of relevant POD significance thresholds on XtX to identify genomic regions harboring footprints of selection. To that end, the UMD3.1 bovine genome assembly (Liu *et al.* 2009) was first split into 5,400 consecutive 1-Mb windows (with a 500 kb overlap). Windows with at least two SNPs displaying XtX> 35.4 (the 0.1% POD threshold) were deemed significant and overlapping “significant” windows were further merged to delineate significant regions. Among the 15 resulting regions, two regions were discarded because their peak XtX value was lower than 40.0 (the 0.01% POD threshold). As detailed in Table 4, the 13 remaining regions lie within or overlap with a Core Selective Sweep (CSS) as defined in the recent meta-analysis by Gutiérrez-Gil *et al.* (2015). This study combined results of 21 published genome-scans performed on European cattle populations using various alternative approaches. The proximity of the XtX peak allows to define positional candidate genes (Table 4) that have, for most regions, already been proposed (or demonstrated) to be either under selection or to control genes involved in traits targeted by selection (see Discussion).

**Table 4.** Regions harboring footprints of selection based on the XtX measure of differentiation and association of the underlying SNPs with SMS (morphology related trait) and piebald coloration differences across the 18 French cattle breeds. For each region, the table gives the peak XtX value (and position in Mb) and the peak BF_is_ and BF_mc_ values in dB units (and positions in Mb) for each traits if the evidence for association is decisive (n.s. if BF < 20). The Table also gives the overlapping Core Selective Sweeps (CSS) regions (with their corresponding sizes and the number of supporting studies) from the meta-analysis by Gutiérrez-Gil *et al.* (2015). Finally, putative underlying candidate genes (and associated candidate functions) are proposed (see the main text).

| ID | Region<br>size in Mb | Overlapping CSS<sup>a</sup><br>size in Mb (#studies) | XtX<br>(Peak Position) | BF<sub>is</sub>-BF<sub>mc</sub><br>for morphology | BF<sub>is</sub>-BF<sub>mc</sub><br>for piebald | Candidate Gene<br>(Function) |
| --- | --- | --- | --- | --- | --- | --- |
| #1 | BTA02:4.17-8.64<br>4.47 | CSS-32<br>13.8 (8) | 58.4<br>(6.70) | n.s.-n.s. | n.s.-n.s. | MSTN:6.214-6.220<br>(conformation) |
| #2 | BTA04:76.7-78.6<br>1.93 | CSS-93<br>2.17 (4) | 45.8<br>(77.6) | n.s.-n.s. | n.s.-n.s. | NUDCD3: 77.599-77.670<br>(unknown) |
| #3 | BTA05:18.0-19.5<br>1.50 | CSS-103<br>1.77 (2) | 76.3<br>(18.5) | n.s.-n.s. | 69.04-52.96<br>(18.5) | KITLG:18.318-18.377<br>(pigmentation) |
| #4 | BTA05:54.7-58.6<br>3.93 | CSS-109<br>22.3 (9) | 54.7<br>(57.6) | 26.6-n.s.<br>(57.6) | n.s.-n.s. | RPS26:57.604-57.607<br>(unknown) |
| #5 | BTA06:17.6-19.2<br>1.54 | CSS-117<br>0.01 (1) | 63.2<br>(18.2) | n.s.-n.s. | n.s.-n.s. | LEF1:18.335-18.451<br>(pigmentation) |
| #6 | BTA06:37.8-40.2<br>2.40 | CSS-123<br>5.09 (8) | 69.4<br>(38.6) | n.s.-n.s. | n.s.-n.s. | LAP3:38.575-38.600<br>(conformation/dairy traits) |
| #7 | BTA06:65.5-74.9<br>9.38 | CSS-130<br>15.3 (12) | 55.6<br>(72.5) | n.s.-n.s.<br>n.s.-n.s. | 37.42-26.45<br>(71.9) | KIT:71.796-71.917<br>(pigmentation) |
| #8 | BTA06:89.6-90.6<br>1.02 | CSS-130<br>13.3 (3) | 40.7<br>(90.2) | n.s.-n.s.<br>n.s.-n.s. | 52.07-38.76<br>(90.2) | ALB:90.233-90.251<br>(pigmentation?) |
| #9 | BTA07:46.4-47.8<br>1.48 | CSS-141<br>12.5 (10) | 46.5<br>(47.3) | n.s.-n.s. | n.s.-n.s. | VDAC1:47.248-47.273<br>(reproduction?) |
| #10 | BTA08:61.4-63.3<br>1.94 | CSS-162<br>0.06 (1) | 49.8<br>(61.8) | n.s.-n.s.<br>n.s.-n.s. | n.s.-n.s.<br>n.s.-n.s. | PAX5:61.400-61.580<br>(pigmentation) |
| #11 | BTA13:56.6-58.6<br>1.98 | CSS-248<br>10.4 (5) | 71.6<br>(57.5) | 23.7-n.s.<br>(58.5) | 26.58-n.s.<br>(57.6) | EDN3:57.571-57.597<br>(pigmentation) |
| #12 | BTA14:22.1-28.8<br>6.76 | CSS-254<br>7.96 (7) | 52.0<br>(24.4) | 35.7-n.s.<br>(24.6) | n.s.-n.s. | PLAG1:25.007-25.009<br>(conformation) |
| #13 | BTA18:13.3-16.0<br>2.75 | CSS-297<br>14.2 (10) | 51.8<br>(14.5) | n.s.-n.s. | n.s.-n.s. | MC1R:14.757-14.759<br>(pigmentation) |

<sup>a</sup> Full descriptions of the CSS (including references to the original studies) are provided in Table S2 by Gutiérrez-Gil et al. \(2015\)

To illustrate how information provided by population-specific covariables might help in formulating or even testing hypotheses to explain the origin of the observed footprints of selection, characteristics of the 18 cattle populations for traits related to morphology (SMS) and coat pigmentation (piebald pattern) were further analyzed within the framework developed in this study. An across population genome–wide association studies was thus carried out under both the STD and AUX models (with 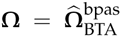) allowing the computation for each SNP of the corresponding BF_is_ and BF_mc_ estimates (Figure S12), and eBP_is_ and eBP_mc_ estimates (Figure S13). We hereafter concentrated on results obtained with BF which are more grounded from a decision theory point of view (and roughly lead to similar conclusions than eBP). For both traits, the BF_is_ resulted in larger BF estimates and a higher number of significant association signals (e.g., at the 20 dB threshold) than BF_mc_. This trend was confirmed by analyzing the POD. Indeed, the 99.9% BF_is_ (respectively BF_mc_) quantiles were equal to 24.9 dB (respectively 18.3 dB) for association with SMS and to 26.3 dB (respectively 11.7 dB) for association with piebald pattern. Nevertheless, at the BF threshold of 20 dB, the false discovery rate for BF_is_ remained small (0.035%) and similar to the one obtained on the simulation studies (e.g., Table 2). Interestingly, among the 13 regions identified in Table 4, three contained (regions #4, #11 and #12) at least one SNP significantly associated with SMS based on the BF_is_ > 20 criterion and none with BF_mc_ > 20 criterion (although BF_mc_ > 5 for the peak of region #12 providing substantial evidence according to the Jeffreys’ rule). For the piebald pattern, results were more consistent since out of the four regions (regions #3, #7, #8 and #11) that contained at least one SNP with a BF_is_ > 20, the BF_mc_ of the corresponding peak SNP was also > 20 (although lower) for all but region #11 (although BF_mc_ = 14.4 for the peak providing strong evidence according to the Jeffreys’ rule). Except for region #7 where both BF peaks lied within the KIT gene (and to a lesser extent for region #11 with SMS), the BF peaks colocalized with (regions #3, #4 and #8) or were very close (less than 50kb) to the XtX peaks. Accordingly, the corresponding XtX estimates decreased when estimated under the STD model, i.e. accounting for the covariables (Figure S14). For instance, the SNP under the XtX peak dropped from 76.3 to 50.3 (from 40.7 to 19.3) for region #3 (respectively region #8). Overall, the posterior means of the individual SNP *β_i_* regression coefficients estimated under the STD model ranged (in absolute value) from 2.2 × 10^−6^ (respectively 1.0 × 10^−8^) to 0.166 (respectively 0.233) for SMS (respectively piebald pattern). These estimates remained close to those derived from Importance Sampling algorithm, although the latter tended to be lower in absolute value (Figure S15). As expected from the above simulation studies, estimates obtained under the AUX model tended to be shrunk towards 0, which was particularly striking in the case of SMS (Figure S15).

**Figure 7.**
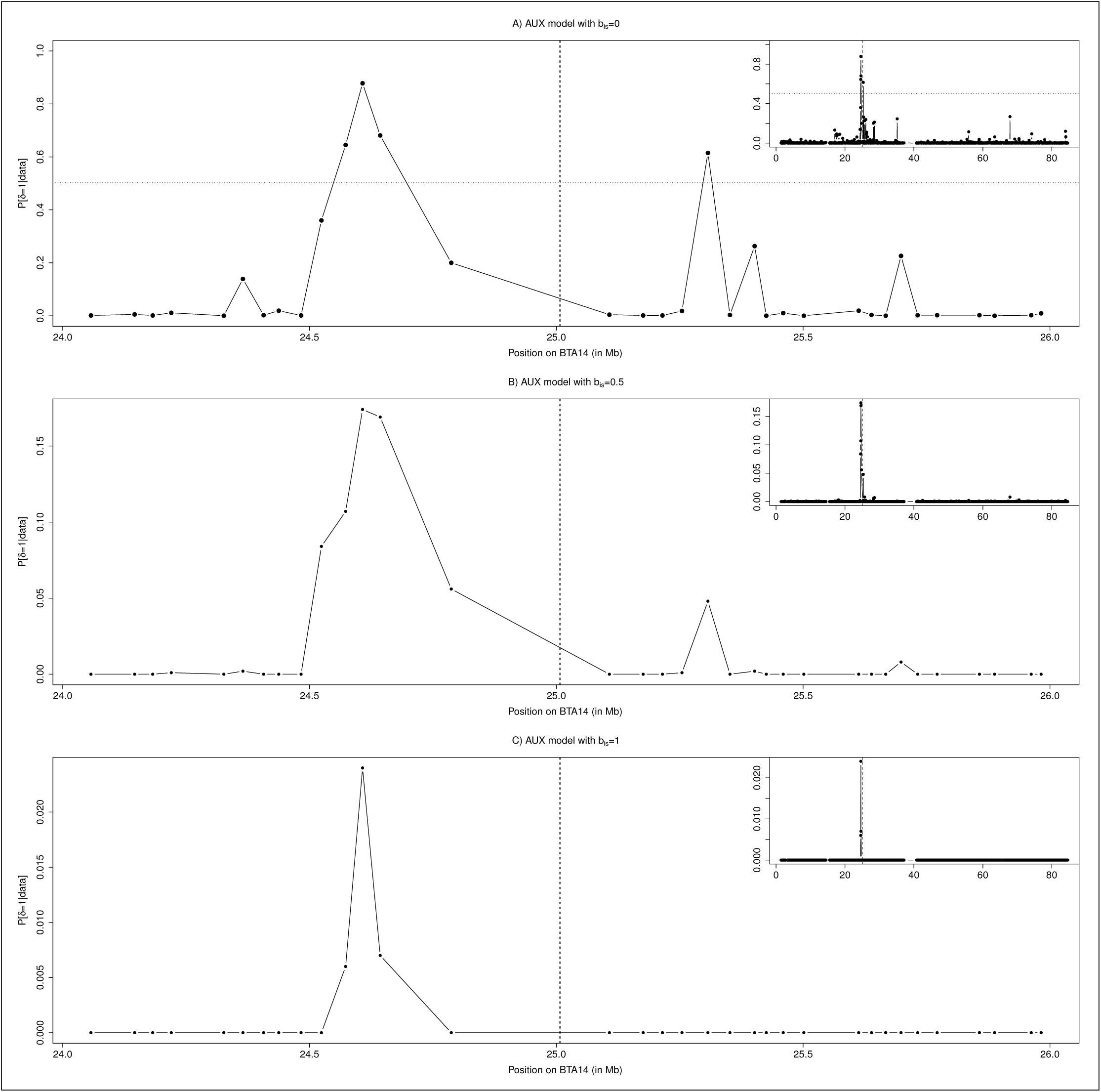
Results of the BTA14 chromosome-wide association analyses with SMS under three different Ising prior parameterizations of the AUX model (A) b_is_ = 0; B) b_is_ = 0.5 and; C) b_is_ = 1.). Plots give, for each SNP, the posterior probability of being associated (P [*δ_i_* = 1 | data]) according to their physical position on the chromosome. The main figure focuses on the region surrounding the candidate gene PLAG1 (positioned on the vertical dotted line) while results over the whole chromosome are represented in the upper left corner. In A), the horizontal dotted line represents the threshold for decisive evidence (corresponding to *BF* = 20 dB).

Finally, analyses of association with SMS were conducted under the AUX model with three different Ising prior parameterizations (b_is_ = 0, b_is_ = 0.5 and b_is_ = 1.) focusing on the 1,394 SNPs mapping to BTA14 (Figure 7). Under the b_is_ = 0 parametrization (equivalent to the AUX model analysis conducted above on a whole genome basis), four SNPs (all lying within region #12) displayed significant signals of association at the *BF* = 20 dB threshold with a peak BF_mc_ value of 28.5 dB at position 24.6 Mb (Figure 7A). These results, obtained on a chromosome-wide basis, provide additional support to the region #12 signal previously observed. They alternatively suggest that power of the BF_mc_ as computed on a whole genome basis might have been altered by the small proportion of SNPs strongly associated to SMS due to multiple testing issues (which BF_is_ computation is not accounted for). Hence, for SNPs mapping to BTA14, the BF_is_ estimated on the initial genome-wide analysis were almost identical to the BF_is_ (*r* = 0.993) and highly correlated to the BF_mc_ (*r* = 0.805) estimated in the chromosome-wide analysis. As expected from simulation results, increasing is*_β_* lead to refine the position of the peak toward a single SNP mapping about 400 kb upstream the PLAG1 gene (Figure 7B and C).

### **Analysis of the** Littorina saxatilis **Pool–Seq data**

The LSA_ps_ Pool–Seq data set was first analyzed under the core model (with *ρ* = 1). In agreement with previous results (Westram *et al.* 2014), the resulting estimate of the population covariance matrix Ω confirmed that the 12 different Littorina populations cluster at the higher level by geographical location and then by ecotype and replicate (Figure 8A). This analysis also allowed to estimate the XtX for each of the 53,387 SNPs that were further calibrated by analyzing a POD containing 100,000 simulated SNPs to identify outlier SNPs (Figure 8). As for the cattle data analysis, the estimate of Ω on the POD was close to the matrix estimated on the original LSA_ps_ data set (*FMD* = 0.516) although the posterior means of *a_π_* and *b_π_* were slightly higher (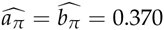 compared to 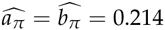 with the LSA_ps_ data set). In total, 169 SNPs subjected to adaptive divergence were found at the 0.01% POD significance threshold. To illustrate how the BayPass models may help discriminating between parallel phenotype divergence from local adaptation, analyses of association were further conducted with ecotype (crab *vs* wave) as a categorical population–specific covariable. Among the 169 XtX outlier SNPs, 65 (respectively 75) displayed BF_is_ > 20 dB (respectively BF_mc_ > 20 dB) (Figure 8B). The two BF estimates resulted in consistent decisions (113 SNPs displaying both BF_mc_ > 20 dB and BF_mc_ > 20 dB), although at the 20 dB more SNPs were found significantly associated under the AUX model (n=176) than with BF_is_ (n=117). Interestingly, several overly differentiated SNPs (high XtX value) were clearly not associated to the population ecotype covariable (small BF). These might thus be responding to other selective pressures (local adaptation) but might also, for some of them, map to sex–chromosomes (Gautier 2014). As a consequence, SNP XtX estimated under the AUX model (i.e., corrected for the “fixed” ecotype effect) remained highly correlated with the XtX estimated under the core model (including for some XtX outliers) with the noticeable exception of the SNPs significantly associated to the ecotype. For the latter, the corrected XtX dropped to values generally far smaller than the 0.01% POD threshold (Figure 8C). Finally, Figure 8D gives the posterior mean of the SNP regression coefficients quantifying the strength of the association with the ecotype covariable. It shows that several SNPs displayed strong association signals 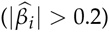 pointing towards candidate genes underlying parallel phenotype divergence. As observed above in the simulation study and in the analysis of the cattle data set, the AUX model estimates tended to be shrunk towards 0, except for the highest values (corresponding to SNPs significantly associated to the covariable) when compared to the estimates obtained under the STD model (Figure S16A). A similar trend for the *β_i_* estimates of the strongly associated SNPs was observed with the importance sampling estimates (Figure S16B).

**Figure 8.**
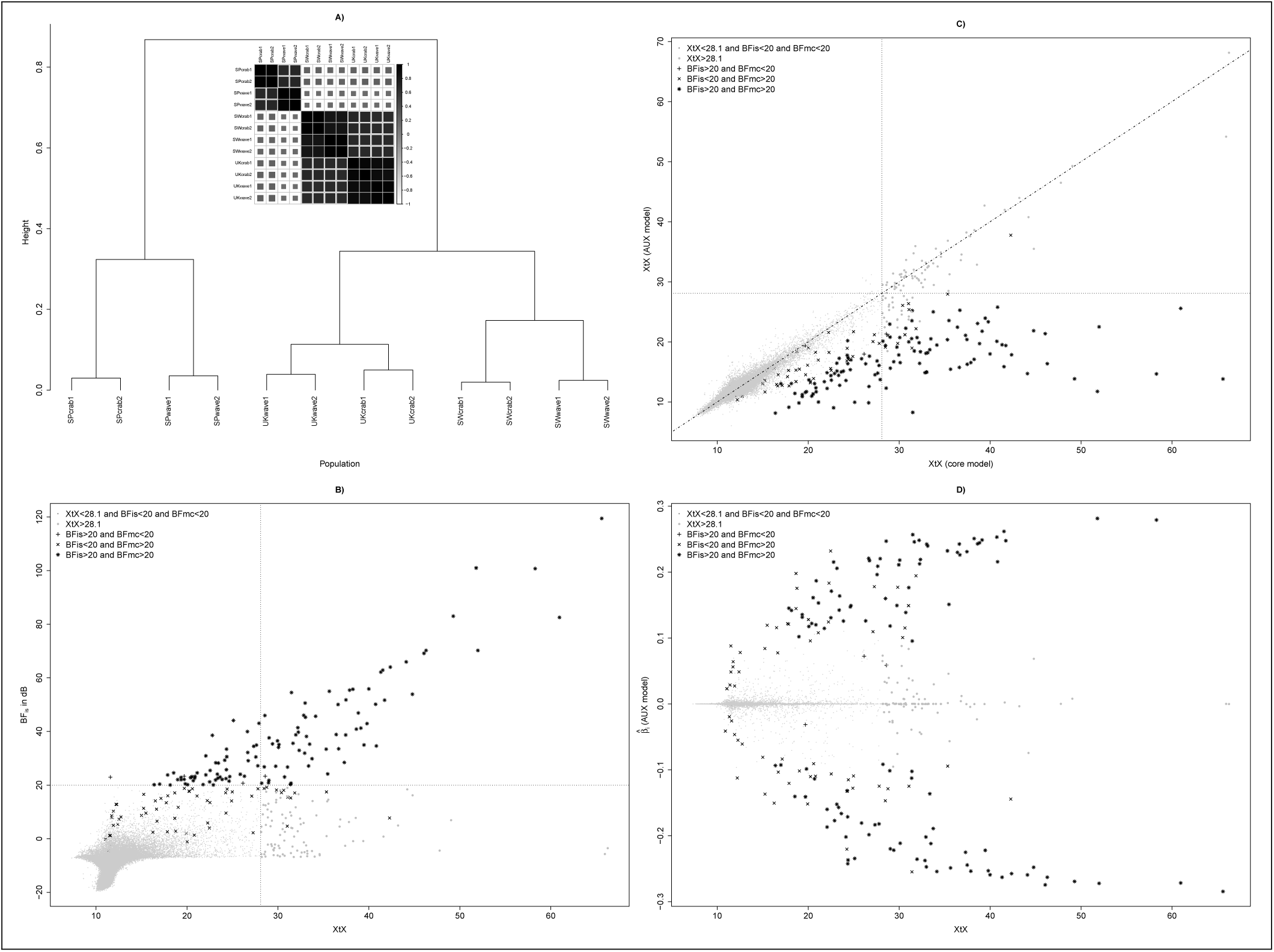
Analysis of the LSA_ps_ Pool–Seq data. A) Inferred relationship among the 12 Littorina populations represented by a correlation plot and a hierarchical clustering tree derived from the matrix Ω estimated under the core model (with *ρ* = 1). Each population code indicates its geographical origin (SP for Spain, SW for Sweden and UK for the United Kingdom), its ecotype (crab or wave) and the replicate number (1 or 2). B) SNP XtX (estimated under the core model) as a function of the BF_is_ for association with the ecotype population covariable. The vertical dotted line represents the 0.1% POD significance threshold (XtX=28.1) and the horizontal dotted line represents the 20 dB threshold for BF. The point symbol indicates significance of the different XtX values, BF_is_ and BF_mc_ (AUX model) estimates. C) SNP XtX corrected for the ecotype population covariable (estimated under the STD model) as a function of XtX estimated under the core model. The vertical and horizontal dotted lines represent the 0.1% POD significance threshold (XtX=28.1). Point symbols follow the same nomenclature as in B). D) Estimates of SNP regression coefficients (*β<u>_i_</u>*) on the ecotype population covariable (under the AUX model) as a function of XtX. Point symbols follow the same nomenclature as in B).

## Discussion

The main purpose of this study was to develop a general and robust Bayesian framework to identify genomic regions subjected to adaptive divergence across populations by extending the approach first described in Coop *et al.* (2010) and Günther and Coop (2013). Because of the central role played in the underlying models by the scaled population covariance matrix (Ω), a first objective was to improve the precision of its estimation. To that end, instead of defining an Inverse-Wishart prior on Ω as in Coop *et al.* (2010), a Wishart prior defined on the precision matrix Λ (Λ = Ω^−1^) was rather considered and equivalently parametrized with an identity scale matrix but varying number of degrees of freedom (*ρ*). As the extensive simulation study revealed, the most accurate estimates were obtained by setting *ρ* = 1 (instead of the number of populations which is equivalent to Coop *et al.* (2010)) leading to a weaker (and singular) informative Wishart prior. Although flexible, the purely instrumental nature of the Ω prior parametrization considered in our models makes it difficult to incorporate prior and possibly relevant information about the populations under study. For instance, spatially (Guillot *et al.* 2014) or even phylogenetically explicit prior might represent in some context attractive alternatives, borrowing for the latter on population genetics theory to model the effect of the demographic history on the covariance matrix (Lipson *et al.* 2013; Pickrell and Pritchard 2012). Apart from investigating different Ω prior specification, additional levels in the hierarchical models were also introduced to estimate the parameters of the (Beta) prior distribution on the ancestral allele frequency. Interestingly, estimating these parameters improved robustness to the SNP ascertainment scheme, in particular when the allele frequency spectrum is biased towards poorly informative SNPs as generally obtained with data from whole genome sequencing experiment (e.g., Pool–Seq data). Simulation results on MAF filtered data sets also suggested that these additional levels might reduce sensitivity of the models to SNP ascertainment bias characterizing genotyping data obtained from SNP chip. Finally, inclusion of a moderate proportion of SNPs under selection did not significantly affect estimation of Ω. Overall, it can be concluded that the core model parametrized with a weakly informative Wishart prior (*ρ* = 1) and that includes the estimation of the parameters *a_π_* and *b_π_* provides a general robust and accurate approach to estimate Ω even with a few thousands of genotyped SNPs. It should also be noted that it outperforms previous implementations carried out under a similar hierarchical Bayesian framework, as in the BayEnv2 software (Coop *et al.* 2010), or relying on moment-based estimators Lipson *et al.* (2013); Bonhomme *et al.* (2010); Pickrell and Pritchard (2012) (see, e.g., Figure 3). As the latter are based on sample allele frequencies, they also remain more sensitive to sample size (and coverage for Pool–Seq data) and, more importantly, they do not allow combining estimation of both the ancestral allele frequencies and covariance matrix that represent a serious issue for small and/or unbalanced designs. Finally, as briefly sketched with visualizations based on correlation plot or hierarchical trees in the present study, the estimation procedure implemented in the BayPass core model might be quite relevant for demographic inference purposes since the matrix Ω has already been shown to be informative about the population history Lipson *et al.* (2013); Pickrell and Pritchard (2012).

Accounting for Ω renders the identification of SNPs subjected to selection less sensitive to the confounding effect of demography (Bonhomme *et al.* 2010; Günther and Coop 2013). To that end the XtX introduced by Günther and Coop (2013) provides a valuable differentiation measure for genome scan of adaptive divergence. While XtX might be viewed as a Bayesian counterpart of the FLK statistic (Bonhomme *et al.* 2010), its computation allows considering population histories more complex than bifurcating trees (i.e., including migration or ancestral admixture events) not to mention improved precision in the estimation of the underlying Ω. For practical purposes however, defining significance threshold for the XtX remains challenging. Indeed, although the XtX are expected under the neutral model to be *χ*-squared distributed (Günther and Coop 2013), the Bayesian (hierarchical) model based procedure leads to shrink the XtX posterior mean towards their prior mean (Gelman *et al.* 2003). As a consequence, an empirical posterior checking procedure, similar in essence to the one previously used in a similar context (Vitalis *et al.* 2014), was evaluated here. It represents a relevant alternative to an arbitrary threshold although it comes at a cost of some additional computational burden. The procedure indeed consists in analyzing (POD) data simulated under the inference model with hyperparameters Ω, *a_π_* and *b_π_* set equal to those estimated on the real data. Comparing the Ω, *a_π_* and *b_π_* estimates obtained on the POD to the original ones ensures that the simulated data provide good surrogates to neutrally evolving SNPs under a demographic history similar to that of the sampled populations. More generally, given the efficiency of the simulation procedure, such simulated data sets might also be relevant to investigate the properties of other estimators of genetic diversity or to evaluate the robustness of various approaches to demographic confounding factors. In the context of this study, a better estimation of Ω was hence shown to improve the performance of the XtX-based differentiation test and association studies with population–specific covariables under the STD and AUX covariate models.

Based on the STD model, Coop *et al.* (2010) relied on an importance sampling (*BF*_is_) estimates of the BF to assess association of allele frequency differences with population–specific covariables. A major advantage of this algorithm stems from its computational efficiency, since only parameter samples drawn from the core model are required. However, the simulation study showed that estimating the *β_i_* regression coefficients with this approach tended to bias (sometimes strongly) the estimates towards zero, as opposed to the posterior means from MCMC parameters values sampled under the STD model. Accordingly, the performances of decision criteria based on eBP’s, that measure to which extent the posterior distribution of the *β_i_* departs from 0, were generally poorer for the *eBP*_is_ than *BF*_mc_. In addition, while a POD calibration, similar to the XtX one considered above is straightforward to apply in practice, eBP (*eBP*_is_ and *eBP*_mc_) and *BF*_is_ could not *per se* deal with multiple testing issues. As previously proposed in a similar modeling context (Riebler *et al.* 2008), introducing binary auxiliary variables attached to each SNP to indicate whether or not they are associated to a given population covariable allows to circumvent these limitations. The resulting *BF*_mc_ showed indeed improved power at stringent decision threshold in the simulation study under the inference model compared to *BF*_is_. In analyses of real data sets, whereas *BF*_is_ estimates were found similar to the *BF*_mc_ ones in analysis of association with ecotype in the Littorina data set, they lead to inflated estimates with the cattle data and thus more (possibly false) significant signals. As shown by analyses on data sets simulated in more realistic scenarios, the intrinsic multiple testing correction (through the prior on the auxiliary variable) might in turn affect the power of *BF*_mc_ decision based criterion. This might explain differences between the results obtained with genome-wide and chromosome-wide analyses of association with the morphology trait in cattle for the region surrounding the PLAG1 gene (BTA14). Besides, in the context of dense genomic data, the AUX model might also be viewed as relevant to more focused analyses for validation (e.g., of genome–wide *BF*_is_ signals) and fine mapping purposes. Hence, the Ising prior on the SNP auxiliary variable provides a straightforward and computationally efficient modeling option to account for the spatial dependency among the neighboring markers (Duforet-Frebourg *et al.* 2014). Prior definition of the b_is_ parameter represents however a first limitation of the AUX model, as defined in this study, and estimating it via an additional hierarchical level would be computationally demanding due to the handling of the normalizing constant (e.g., Marin and Robert 2014, ch. 8.3). Comparing the results from different analyses with increasing values of b_is_ thus appears as a valuable empirical strategy. More importantly, it should also be noticed that the Ising prior essentially consists in a local smoothing of the association signals whose similarity stems from a correlation of the underlying allele frequencies (across all the populations). It thus does not fully capture LD information contained in the local haplotype structure. To that end further extensions of the AUX (and STD) model following the hapFLK method (Fariello *et al.* 2013) that directly relies on haplotype information, might be particularly appropriate although difficult to envision for data originating from Pool–Seq experiments.

As expected, in both simulated and real data sets, SNPs strongly associated (|*β_i_*|>0.2) with a given covariable tended to be overly differentiated (high XtX value). Interestingly however, the STD and AUX covariate models remained more powerful to identify SNPs displaying weaker association signal (typically with |*β_i_*| < 0.1) for which the XtX values did not overly depart from that of neutral SNPs. Providing information on an underlying covariable (or a proxy of it) is available, the STD and AUX models might thus allow to identify SNPs within soft adaptive sweeps or subjected to polygenic adaptation, these types of selection schemes leading to more subtle population allele frequencies differences difficult to detect (e.g., Pritchard *et al.* 2010). Conversely, the covariate models were shown to correct the XtX differentiation measure for the fixed effects of the considered population–specific covariables, refining the biological interpretation of the remaining overly differentiated SNPs by excluding these covariable as key drivers. In principle, across–population association analyses could be performed with any population–specific covariable like environmental covariables (Coop *et al.* 2010; Günther and Coop 2013) but also categorical or quantitative traits as illustrated in examples treated in this study. As such, the STD and AUX covariate models might also be viewed as powerful alternatives to Q_ST_–F_ST_ comparisons to assess divergence of quantitative traits (see Leinonen *et al.* 2013, for review) by accurately incorporating genomic information to account for the neutral covariance structure across population allele frequencies. Yet, it should be kept in mind that the considered models only capture linear relationships between allele frequencies differences and the covariable. Apart from possibly lacking power for more complex types of dependency, the correlative (and not causative) nature of the association signals might be misleading, noticeably when the (unobserved) causal covariable is correlated with the analyzed trait or with the principal axes of the covariance matrix (Günther and Coop 2013). Nevertheless, increasing the number of populations and (if possible) the number of studied covariables should overcome these limitations. Still, when jointly considering several covariables, this also advocates for an orthogonal transformation (and scaling) step, e.g. using PCA, to better assess their relationships and to further perform analysis of association on an uncorrelated set of covariables (e.g., principal components).

As a proof of concept, analyses were carried out on real data sets from both model and non model species. Results obtained for the French cattle data demonstrated the versatility of the approach and illustrated how association studies could give insights into the putative selective forces targeting footprints of selection. As a matter of expedience we only hereby focused on the thirteen strongest differentiation signals. As expected from the importance of coat pigmentation in the definition of breed standards, at least six genomic regions contained genes known to be associated to coat color and patterning variation, in agreement with previous genome scan for footprints of selection (see Gutiérrez-Gil *et al.* 2015, for review). These include MC1R (region #13) that corresponds to the locus *Extension* with three alleles identified to date in cattle responsible for the red, black (or combination of both) colors (Seo *et al.* 2007). Similarly, variants localized within the KIT (region #7) and PAX5 (region #10) genes were found highly associated to patterned pigmentation (proportion of black) in Hosltein, accounting for respectively 9.4% and 6.0% of the trait variance (Hayes *et al.* 2010). Within region #7, KIT clusters with KDR (closest to the XtX peak) and PDGFRA, two other Tyrosine kinase receptor genes that have also been proposed as candidate coloration genes under selection in other studies (Flori *et al.* 2009; Gutiérrez-Gil *et al.* 2015; Qanbari *et al.* 2014). In region #11, the XtX peak was less than 25 kb upstream EDN3 that is involved in melanocyte development and within which mutations were found associated to pigmentation defects in mouse, human and also chicken (Bennett and Lamoreux 2003; Dorshorst *et al.* 2011; Saldana-Caboverde and Kos 2010). Accordingly, Qanbari *et al.* (2014) recently found a variant in the vicinity of EDN3 strongly associated with coat spotting phenotype of bulls (measured as the proportion of their daughters without spotting) in the Fleckvieh breed. The peak in region #2 was 100 kb upstream the KITLG gene which is involved in the roan phenotype (mixture of pigmented and white hairs) observed in several cattle breeds (Seitz *et al.* 1999). Mutations in this gene have also been found to underlie skin pigmentation diseases in human (Picardo and Cardinali 2011). Finally, region #5 contains the LEF1 gene (100 kb from the XtX peak) that has recently been demonstrated to be tightly involved in blond hair color in (human) Europeans (Guenther *et al.* 2014). Three other regions contained genes that affect cattle body conformation. These include region #1 containing the myostatin gene (MSTN), one of the best known examples of economically important genes in farm animals since it plays an inhibitory role in the development and regulation of skeletal muscle mass (Stinckens *et al.* 2011). MSTN is in particular responsible for the so-called double muscling phenotype in cattle (Grobet *et al.* 1997). Region #12 contains PLAG1 that has been demonstrated to influence bovine stature (Karim *et al.* 2011). Similarly, region #6 contain encompasses the NCAPG-LCORL cluster in which several polymorphisms have been found strongly associated to height in human (Allen *et al.* 2010), horse (Signer-Hasler *et al.* 2012) and cattle (Pryce *et al.* 2011). However, combining results from a genome-scan for adaptive selection with a comprehensive genome-wide association study with milk production traits in the Holstein cattle breed, Xu *et al.* (2015) proposed the LAP3 gene (within which the XtX peak mapped) as the main driver of a selective sweep overlapping with region #12. Regarding the four remaining regions (#2,#4,#8 and #9), the retained candidate genes corresponded to the gene within which the XtX peak is located (NUDCD3, RPS26 and VDAC1 for regions #2, #4 and #9 respectively) or is the closest (less than 15 kb from ALB for region #8). As for RPS26, although NUDCD3 has been highlighted in other studies (e.g., Flori *et al.* 2009; Xu *et al.* 2015), the poorly known function of these genes makes highly speculative any interpretation of the origin of the signals. Conversely, the various and important roles played by ALB (bovine serum albumin precursor) do not allow a clear hypothesis to be formulated about the trait underlying the region #8 signal. More presumably, due to the role of VDAC1 in male fertility (Kwon *et al.* 2013), the footprint of selection observed in region #9 might result from selection for a trait related to reproduction. Overall, association analyses carried out under the covariate models revealed strong association of SNPs within KITLG (region #3), KIT (region #7) and EDN3 (region #11) with variation in the piebald pattern across the populations thereby supporting the hypothesis of selection on coat coloration to be the main driver of the three corresponding signatures of selection. These results also confirm the already well known key role of these genes on coloration patterning. Interestingly, the observed association signals within ALB (region #8) also suggest that this gene might influence coat coloration in cattle which, to our knowledge, has not been previously reported. Finally, association studies on the SMS trait suggested that PLAG1 (region #12) has been under strong selection in European cattle and contribute to morphological differences across the breeds. Yet, the strongest association signals was 400 kb upstream PLAG1 suggesting the existence of some functional variants (possibly in regulatory regions) different from those already reported (Karim *et al.* 2011) although such results need to be confirmed with denser SNP data sets. Conversely, no association signal was found within the selection signature under region #6 adding more credits to selection for milk production (Xu *et al.* 2015) as the main underlying adaptive constraint rather than morphological trait as previously hypothesized (see above). Analysis of the *Littorina saxatilis* Pool–Seq data (Westram *et al.* 2014) illustrate how BayPass can be helpful to realize a typology of the markers relative to an ecological covariable in a non–model species. In agreement with the original results, several genes represent good candidate to underlie parallel phenotypic divergence in this organism and might deserve follow-up validation studies. From a practical point of view however, compared to combining several pairwise F_ST_ population tests (Westram *et al.* 2014), the approach proposed here greatly simplified the analyses and the biological interpretation of the results while allowing both an optimal use of the data and a better control for multiple testing issues.

Overall, the models described here and implemented in the software package BayPass provide a general and robust framework to better understand the patterns of genetic divergence across populations at the genomic level. They allow i) an accurate estimation of the scaled covariance matrix whose interpretation gives insights into the history of the studied populations; ii) a robust identification of overly differentiated markers by correcting for confounding demographic effects; and iii) robust analyses of association of SNP with population–specific covariables giving in turn insights into the origin of the observed footprints of selection. In practice, when compared to BayEnv2, BayPass lead to a more accurate and robust estimation of the matrix Ω (and the related measures) and thus improved the performances of the different tests. In addition, various program options were developed to investigate the different modeling extensions, including analyses under the STD and AUX models and exploration of the Ising prior parameters to incorporate LD information. More generally, as demonstrated by the analysis of individual-based simulated data sets, the method developed in this study was found to be among the most efficient in terms of power, robustness and computational cost when compared to the other state-of-the-art or recently developed genome scan methods. Moreover, as opposed to most of the currently available approaches, the different decision measures (XtX, eBP and BF) can be computed for both allele (from standard individual genotyping experiments) and read (from Pool–Seq experiments) count data (while also accommodating missing data). Finally, although computation times scale roughly linearly with the data set complexity (number of populations × number of markers), for very large data sets, several strategies might be efficient to reduce computational burden. For instance, because estimation of Ω was found robust to moderate ascertainment bias, one may filter low polymorphic markers (e.g., overall MAF<0.01) since those are not informative for genome scan purposes, and/or consider sub-sampling of the initial data set (e.g., chromosome-wide analyses).

## Acknowledgments

I wish to thank Anja Westram for providing early access to the Littorina data and Pierre de Villemeureuil for providing the polygenic data sets simulated under the HsIMM, IMM and SS scenarios (and information about the underlying Python/SimuPOP scripts). I also wish to thank the two anonymous reviewers for their valuable comments. I am finally grateful to the genotoul bioinformatics platform Toulouse Midi–Pyrenees for providing computing resources. This work was partly funded by the ERA–Net BiodivERsA2013-48 (EXOTIC), with the national funders FRB, ANR, MEDDE, BELSPO, PT-DLR and DFG, part of the 2012–2013 BiodivERsA call for research proposals.

